# Noise Control In Gene Regulatory Networks With Negative Feedback

**DOI:** 10.1101/049502

**Authors:** Michael Hinczewski, D. Thirumalai

**Affiliations:** Department of Physics, Case Western Reserve University, OH 44106; Department of Chemistry, The University of Texas at Austin, TX 78712

## Abstract

Genes and proteins regulate cellular functions through complex circuits of biochemical reactions. Fluctuations in the components of these regulatory networks result in noise that invariably corrupts the signal, possibly compromising function. Here, we create a practical formalism based on ideas introduced by Wiener and Kolmogorov (WK) for filtering noise in engineered communications systems to quantitatively assess the extent to which noise can be controlled in biological processes involving negative feedback. Application of the theory, which reproduces the previously proven scaling of the lower bound for noise suppression in terms of the number of signaling events, shows that a tetracycline repressor-based negative-regulatory gene circuit behaves as a WK filter. For the class of Hill-like nonlinear regulatory functions, this type of filter provides the optimal reduction in noise. Our theoretical approach can be readily combined with experimental measurements of response functions in a wide variety of genetic circuits, to elucidate the general principles by which biological networks minimize noise.

---

The genetic regulatory circuits that control all aspects of life are inherently stochastic. They depend on fluctuating populations of biomolecules interacting across the crowded, thermally agitated interior of the cell. Noise is also exacerbated by low copy numbers of particular proteins and mRNAs, as well as variability in the local environment.^1^^-^^6^ Yet the robust and reproducible functioning of key systems requires mechanisms to filter out fluctuations. For example, regulating noise is relevant in stabilizing cell-fate decisions in embryonic development,^7^ prevention of random switching to proliferating states in cancer-regulating miRNA networks,^8^ and maximization of the efficiency of bacterial chemotaxis along attractant gradients.^9^ Comprehensive analysis of yeast protein expression reveals that proteins involved in translation initiation, ribosome formation, and protein degradation, have lower relative noise levels,^10^ suggesting natural selection could favor noise reduction for certain essential cellular components.^11^,^12^

A common regulatory motif capable of suppressing noise is the negative feedback loop,^1^,^2^,^13^^-^^18^ as has been explicitly demonstrated in synthetic gene circuits.^1^,^14^,^15^ Feedback pathways for a given chemical species can be mediated by numerous signaling molecules, each with its own web of interactions and stochastic characteristics that determine the ultimate effectiveness of the system in damping the fluctuations of the target population and maintaining homeostasis. Thus, uncovering generic laws governing the behavior of such control networks is difficult. A major advance was made by Lestas, Vinnicombe, and Paulsson (LVP),^19^ who showed that information theory can set a rigorous lower bound on the magnitude of fluctuations within an arbitrarily complicated homeostatic negative feedback network. Since the bound scales like the fourth root of the number of signaling events, noise reduction is extremely expensive. This underscores the pervasiveness of biological noise, even in cases where there may be evolutionary pressure to minimize it.

The existence of a rigorous bound raises a number of intriguing issues. Can a biochemical network actually reach this lower bound, and thus optimally suppress fluctuations? What would be the dynamic behavior of such an optimal system, and how would it depend on the noise spectrum of the system components? Here we answer these equations using a theory related to the optimal linear noise-reduction filter, developed by Wiener^20^ and Kolmogorov.^21^ Though the original context of Wiener-Kolmogorov (WK) filter theory was removing noise from corrupted signals in engineered communications systems, it has recently become a powerful tool for characterizing the constraints on signaling in biochemical networks.^22^,^23^ recently, we showed that the action of kinase and phosphatase enzymes on their protein substrates, the basic elements of many cellular signaling pathways, can in fact effectively be represented as an optimal WK filter.^22^ The WK theory also describes how systems like E. coli chemotaxis can optimally anticipate future changes in concentrations of extracellular ligands.^23^ Although the classic WK theory is strictly defined for linear filtering of continuous signals (a reasonable approximation for certain biochemical networks), it can also be extended to yield constraints in the more general case of nonlinear production of molecular species with discrete population values.^22^

Interestingly, for a broad class of systems the WK linear solution turns out to be the global optimum among all nonlinear or linear networks, allowing us to delineate where non-linearity is potentially advantageous in biochemical noise control. Most importantly, since the WK theory is formulated in terms of experimentally accessible dynamic response functions, it also provides a design template for realizing optimality in synthetic circuits. As an illustrative example, we predict that a synthetic autoregulatory TetR loop, engineered in yeast,^24^ can be fine-tuned to approximate an optimal WK filter for TetR mRNA levels. Though a simple design, similar filters could be employed in nature to cope with Poisson noise arising from small copy numbers of mRNAs, often on the order of 10 per cell.^25^ Based on the application of the theory to the synthetic gene network we propose that the extent of noise reduction in biological circuits is determined by competing factors such as functional efficiency, adaptation, and robustness.

## Results

To make the paper readable and as self-contained as possible many of the details of the calculation are relegated to four Appendices. The main text contains only the necessary details needed to follow the results without the distraction of the mathematics.

## Linear response theory for a general control network

To motivate the WK approach for a general control network, we start with the simple case where two species within the network are explicitly singled out:^19^ a target *R* with time-varying population *r(t*) fluctuating around mean 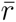, and one of the mediators in the feedback signaling pathway *P*, with population *p(t*) varying around 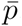. We assume a continuum Langevin description of the dynamics,^13^,^16^,^26^,^27^ where the rate

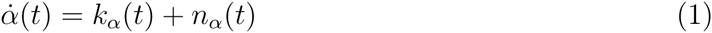

for *α = r*or *p*, can be broken down into deterministic *(κ*_*α*_) and stochastic *(n*_*α*_) parts. The function *κ*_*α*_*(t*) encapsul rates the entire web of biochemical reactions underlying synthesis and degradation of species *α*, and can be an arbitrary functional of the past history of the system up to time *t*. It is typically divided into two parts, 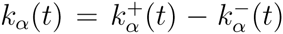, corresponding to the production (+) and destruction (-) rates of the species *α*. The term *n*_*α*_*(t*) is the additive noise contribution, which can also be divided into two parts, 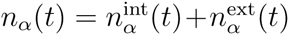. The first is the “intrinsic” or shot noise, arising from the stochastic Poisson nature of *α* generation, 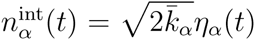, where 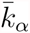 is the mean production rate, or equivalently the mean destruction rate, 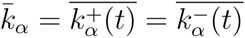, and *η*_*α*_*(t*) is a Gaussian white noise function with correlation 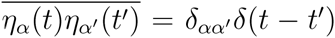. The second part, 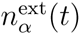, is “extrinsic” noise, which arises due to fluctuations in cellular components affecting the dynamics of and *P* that are not explicitly taken into account in the two-species picture. These could include mediators in the signaling pathway, or global factors likeibosome and RNA polymerase levels. For simplicity, our main focus will be the case of no extrinsic noise. However, we will show later how a straightforward extension of the theory reveals that the same system can behave like an optimal WK filter under a variety of extrinsic noise conditions.

For small deviations *δα(t) = α(t)-κ* from the mean populations *κ, κ*_*α*_*(t*) can be linearized withespect to *δα(t)*,

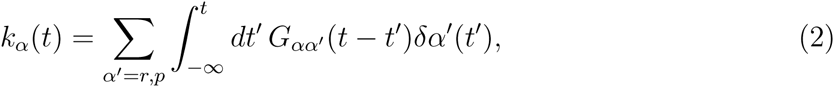

where *G*_*αα*'_*(t*) are linear response functions, which express the dependence of *κ*_*α*_*(t*) on the past history of *δα*'*(t*). The functions *G*_*αα*'_*(t*) capture the essential characteristic responses of the control network to perturbations away from equilibrium (Fig. 1). In the static limit, *G*_*αα*'_*(t*) have appeared in various guises as gains,^6^ susceptibilities,^17^ or steady-state Jacobian matrices,^27^ and in the frequency-domain as loop transfer functions.^13^,^16^ Feedback between *R* and *P* is encoded in the cross-responses *G*_*rp*_*(t*) and *G*_*pr*_*(t*). In the simplest scenario, the only non-zero self-responses *G*_*αα*_*(t*) are decay terms, 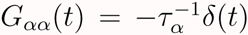, where *τ*_*α*_ is the decay time scale for species *α*. However, the theory works generally for more complicated self-response mechanisms.

**Figure 1:**
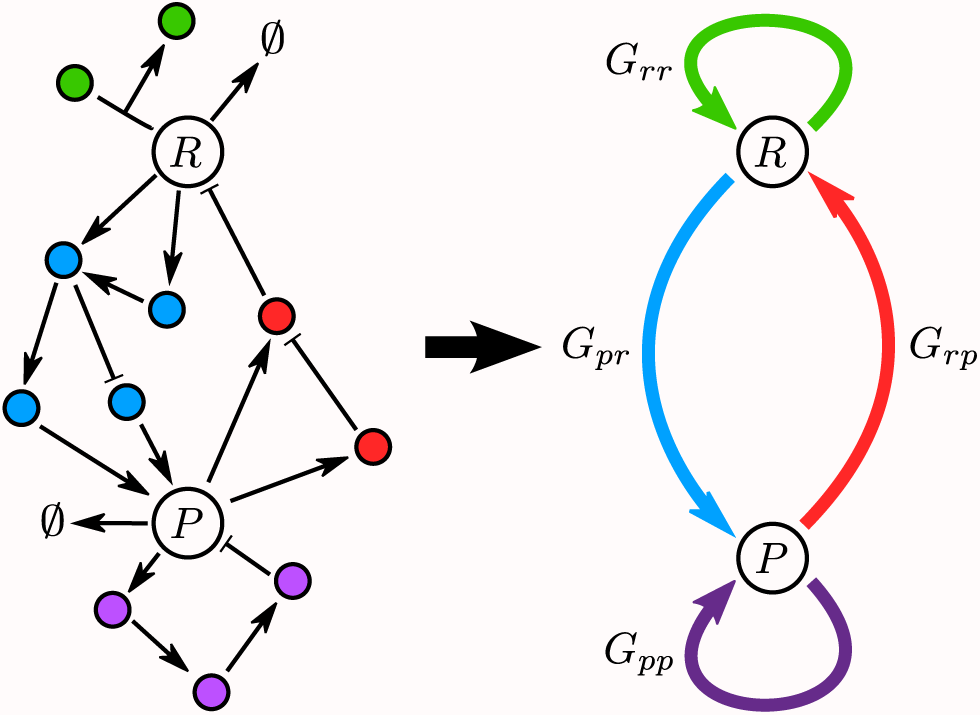
Schematic of a complex signaling network with the target species *R* and one mediator *P* singled out. In focusing on two species, the action of all the other components is effectively encoded in four response functions—*G*_*rr*_*(t*), *G*_*pp*_*(t*), *G*_*rp*_*(t*), *G*_*pr*_*(t*)—that describe how the entire dynamical system responds to perturbations in *R* and *P*.

## Control network as a noise filter

The connection between the linearized dynamical description and WK filter theory arises from comparing the original system to the case where feedback is turned off (i.e. setting *G*_*rp*_*(ω*) or *G*_*pr*_*(ω*) to zero). Let us define a few terms to make the noise filter analogy clear. Without feedback, the target fluctuations are *δr*_0_*(t) ≡ s(t*), where we denote *s(t*) the *signal*. This is to distinguish it from *δr(t*) in the original system, which is the *output*. The difference between the two, which reflects the impact of the feedback network, we express as 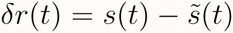, where 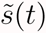 is referred to as the *estimate*. In this analogy, minimizing *δr(t*) requires a feedback loop where the estimate 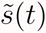 is as close as possible to the signal *s(t*). The only thing left to specify is the relationship between 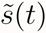 and *s(t*).

The dynamical system in Eqs. (1)-(2) takes a simple form in Fourier space, where the fluctuations *δα(ω*) satisfy:

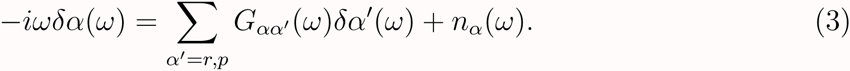

We solve Eq. (3) for *δr(ω*) and break up the *R* fluctuation into two contributions, 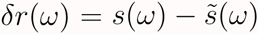, with the signal *s(ω*) and estimate 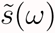 given by:

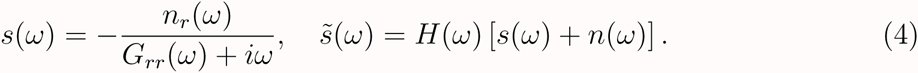

Here we have introduced a noise function *n(ω*),

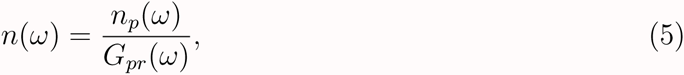

and a filter function *H(ω*):

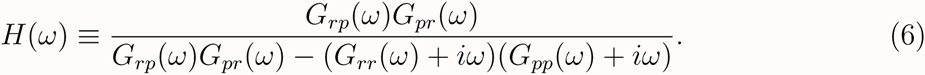

Thus in the time domain the estimate 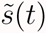 is the convolution of the filter function *H(t*) and a noise-corrupted signal *y(t*) ≡ *s(t*) + *n(t)*,

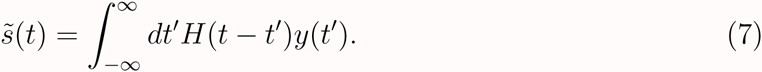

Eqs. (4)-(6) constitute a one-to-one mapping between the linear response and noise filter descriptions of the system in Fourier space. They relate the four filter quantities, *s(ω*), 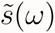, *n(ω*), and *H(ω*), to the four linear response functions *G*_*rr*_*(ω*), *G*_*rp*_*(ω*), *G*_*pr*_*(ω*), and *G*_*pp*_*(ω*).

The entire noise filter system is illustrated schematically in Fig. 2. Note that the noise function in the filter analogy, *n(t*), is related to *n*_*p*_*(t*) in Fourier space as *n(ω*) = *n*_*p*_*(ω*)/*G*_*pr*_*(ω*). Thus, the stochastic nature of the mediator *P* production makes estimation non-trivial, since the function *H(t*) must try to filter out the *n(t*) component in *y(t*) in order to produce 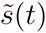 close to *s(t*). Though we confine ourselves throughout this work to the case of a dynamical system with a single target and mediator species, one can easily generalize the entire approach to explicitly include many mediators, which could potentially be involved in a complex signaling pathway. The linearized dynamical system in Eqs. (1)-(2) would still have the same form (with index *α* running over all the species of interest), and the mapping onto the filter problem for the target species would be analogous. The only difference is that *n(ω*) and *H(ω*) would be more complicated functions of the various individual noise terms *n*_*α*_*(ω*) and the response functions *G*_*αα*'_*(ω*) of the mediators. In our reduced, two species description, the action of all the unspecified chemical components is effectively included in the four response functions described above, with their stochastic effects contributing to the extrinsic noise. Fig. 1 shows a schematic of such a reduction. The fine-grained details of the signaling pathways connecting our target *R* and mediator *P*, potentially involving many interacting species, are encoded in *G*_*rr*_, *G*_*pp*_, *G*_*rp*_, and *G*_*pr*_. As an example of how this two-species reduction would work in practice, in Appendix B we treat an important example of a feedback loop involving multiple mediators, representing a signaling cascade in series.

**Figure 2:**
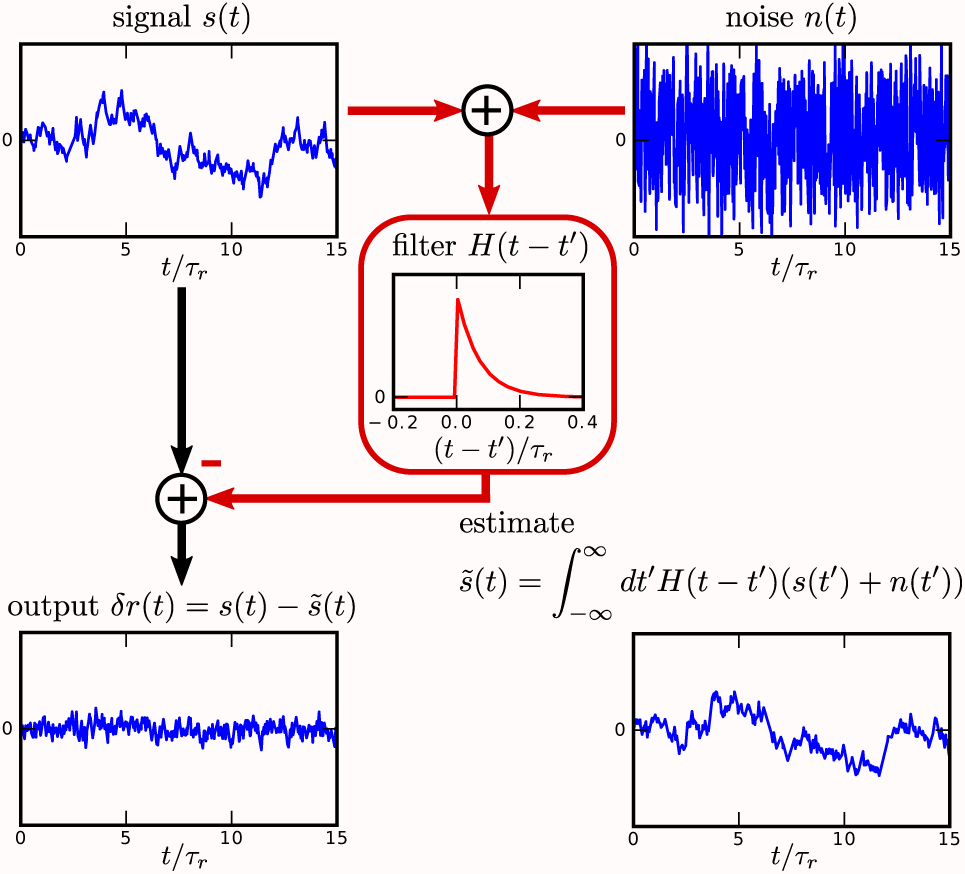
Signal processing diagram illustrating noise suppression in a negative feedback loope-interpreted as a linear filter. The fluctuations in the target species *δr(t*) (lower left) are expressed as 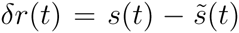, where theaw signal *s(t*) (upper left) equals *δr(t*) in the absence of feedback control, and the estimate 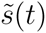 (loweright) is the contribution of the feedback loop. This estimate is given by the convolution of a filter function *H(t*) (center) and the corrupted signal *s(t*) + *n(t*), where *n(t*) is the noise (upperight). The goal of Wiener-Kolmogorov theory is to find a causal *H(t*) such that the standard deviation of *δr(t*) is minimized. All sample trajectories shown in the figure are generated from numerically solving the linearized version of the dynamical system in Eq. (10).

## Wiener-Kolmogorov theory yields the optimal filter

The WK optimization problem consists of minimizing 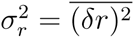, the variance of target fluctuations, which are related to *H(t), s(t*), and *n(t*) through the frequency-domain integral^28^ (see derivation in Appendix A):

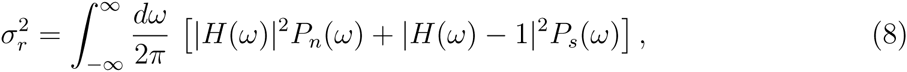

where *H*(*ω*) is the Fourier transform of *H(t*), and *P*_*n*_*(ω*), *P*_*s*_*(ω*) are the power spectral densities (PSD) of *n(t*) and *s(t*) respectively, i.e. the Fourier transforms of their autocorrelation functions. If *P*_*n*_*(ω*) and *P*_*s*_*(ω*) are given, the task is to minimize 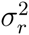 in Eq. (8) over all possible *H(ω*). The main constraint that makes the solution mathematically difficult is that *H(ω*) must correspond to a physically realizable control network, which imposes the crucial restriction that the time-domain convolution function *H(t*) must be causal, depending only on the past history of the input, *H(t*) = 0 for *t<*0. The great achievement of Wiener and Kolmogorov was to derive the form of the optimal causal solution *H*_opt_*(ω*):

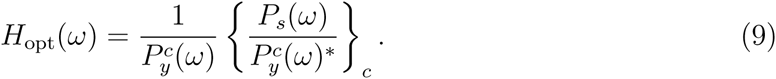

The *c* super/subscripts refer to two different decompositions in the frequency domain which enforce causality: (i) Any physical PSD, in this case *P*_*y*_*(ω*) corresponding to the corrupted signal *y(t) = s(t*) + *n(t*), can be written as 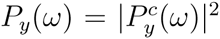. The factor 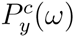, if treated as a function over the complex *ω* plane, contains no zeros and poles in the upper half-plane (Im*ω* > 0).^29^ (ii) We also define an additive decomposition denoted by *{F(ω)}*_*c*_ (see Appendix A) for any function *F(ω*), which consists of all terms in the partial fraction expansion of *F(ω*) with no poles in the upper half-plane. In Appendix A we provide in detail a new derivation of Eq. (9), the heart of the WK theory.

## Optimal noise control in a yeast gene circuit with feedback

To illustrate the nature of the optimal WK solution we choose as a case study the yeast negative autoregulatory gene circuit designed by Nevozhay *et. al.,^24^* drawn schematically in Fig. 3(a). The gene encoding for the TetR protein is under the control of the *P*_GAL1-D12_ promoter, whose activity can be repressed by binding TetR dimers. The strength of the feedback can be modulated by changing the extracellular concentration *A* of the inducer anhydrotetracycline (ATc), which enters the cell, binds to TetR and prevents its association with the promoter, thus weakening repression.

**Figure 3:**
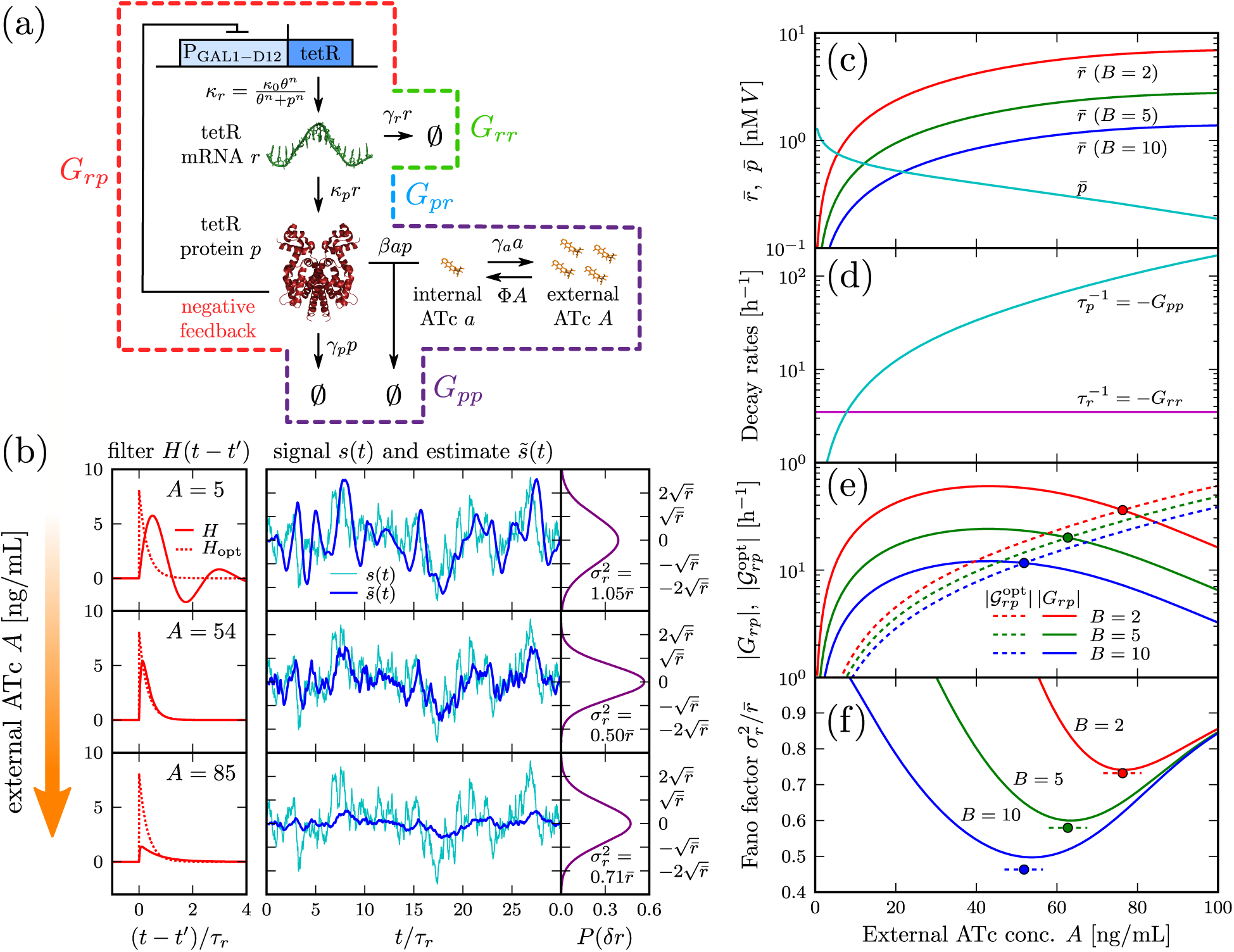
(a) The synthetic yeast gene circuit designed by Nevozhay *et. al*.^24^ The TetR protein negatively regulates itself by binding to its own promoter. The inducer molecule ATc associates with TetR, inhibiting its repressor activity. The subsequent panels show results for this gene circuit using the linear filter theory applied to the dynamical model of Eq. (10), with experimentally-derived parameters (Table 1). (b) Filter functions *H(t*) and *H*_opt_*(t*), sample signal *s(t*) and estimate 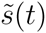 time series for burst ratio *B* = 10 and three different values of extracellular ATc concentration *A* [ng/mL]. *H(t*) is from Eq. (19), while *H*_opt_*(t*) is from Eq. (18). The sample time series trajectories are numerical solutions of the linearized Eq. (10). On the right are the resulting equilibrium probability distributions *P(δr*), where 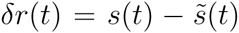, which are Gaussians with variance 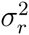. For *A* ≈ 54 ng/mL, the circuit approximately functions as an optimal WK filter *(H(t*) is close to *H*_opt_*(t)*), maximally suppressing fluctuations in the population levels of TetR mRNA (minimizing 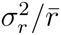). (c) Mean populations of free intracellular TetR mRNA, 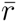, and TetR protein dimers, 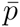. (d) The decay rates of free mRNA and proteins, 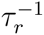 and 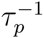, which are related to the network self-response functions *G*_*rr*_ and *G*_*pp*_ (both are constants in the frequency domain as shown in Eq. 11). (e) The magnitude of the network cross-response, |*G*_*rp*_| (solid lines), plotted together with the optimal magnitude 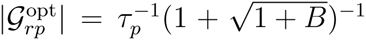 (dashed lines). Filled circles mark the intersection defining *A* = *A*_opt_, where the system behaves approximately like an optimal WK filter. (f) The Fano factor 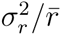 (solid lines), compared to the optimal WK value 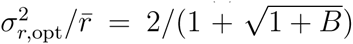 (horizontal dashed lines). Filled circles mark the position *A = A*_opt_.

In order to analyze the TetR negative feedback gene circuit, we start with the simple mathematical model introduced in Ref. 24, which provided results that are consistent with the experimental data. The simplified model, which captures the essence of the synthetic gene network, features as the main variables the population of free intracellular TetR dimer, *p(t*), and free intracellular ATc molecules, *a(t*). In addition to the regulatory loop, the experimental gene circuit has a parallel yEGFP reporter portion, which acts as a monitor of TetR protein levels. Because we focus on the system as a noise filter for the TetR mRNA population, and the yEGFP part does not influence this analysis,^24^ we ignore the reporter circuit.

The production of the TetR dimers occurs in a single step, with the autoregulation of the rate described by a repressory Hill function. We divide this step into two parts, introducing as an additional variable the population of TetR mRNA *r(t*). The feedback loop (Fig. 3(a)) consists of mRNA production at a rate given by the Hill function *κ*_*r*_*(t*) = *κ*_0_*θ*^*n*^/*(θ*^*n*^ + *p*^*n*^*(t*)), followed by TetR dimer generation at a rate given by *κ*_*p*_*r(t*). The degradation/dilution of the mRNA and dimers is modeled through decay terms *γ*_*r*_*r(t*) and *γ*_*p*_*p(t*). We could have modeled additional (comparatively fast) chemical substeps involved in this loop, such as TetR dimerization, the binding of the repressor to the individual promoter sites, or the role of RNAP and ribosomes in the transcription and translation processes. Though we limit ourselves to the two substep description to illustrate the filter theory, the stochastic effects of additional complexity can be approximately treated through general “extrinsic” noise terms incorporated into *n*_*r*_*(t*) and *n*_*p*_*(t*).

The main experimental variable that allows tuning of the yeast gene network behavior is the external ATc concentration *A*, which is assumed to be time independent. As illustrated in Fig. 2(a), there is an influx *ΦA* of ATc molecules into the cell. Once inside, the ATc molecules associate with the TetR at a rate *βa(t)p(t*). Additional loss of intracellular ATc through degradation, outflux, and dilution is modeled through an effective decay rate *γ*_*a*_*a(t*). We assume that the dissociation of ATc from TetR occurs on long enough timescales that it can be ignored. Since the influx/association/outflux of ATc is fast compared to the transcription and translation processes of the main feedback loop, we further assume that *a(t*) instantaneously equilibriates at the current value of *p(t*). Thus, the dependence of *a(t*) on *p(t*) is determined by equating the influx and total loss rate, which leads to *a(p(t)) = ΦA*/*(γ*_*a*_ + *βp(t)*).

For the model described above, the dynamical equations for *r(t*) and *p(t*) are,

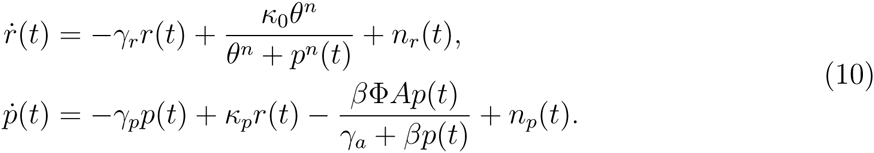

The parameters, with values derived from experimental fitting,^24^ are listed in Table 1. The only quantity that is not independently known from the fit is the rate *κ*_*p*_, which we allow to vary in the range *κ_*p*_/γ*_*r*_ ≡ *B* = 2 - 10, comparable to typical experimentally measured protein burst sizes.^30^ Setting the right sides of Eq. (10) to zero, and averaging over *n*_*r*_*(t*) and *n*_*p*_*(t*), we numerically solve for the equilibrium values 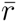 and 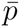 as a function of external ATc concentration *A* [Fig. 3(c)]. For *A* = 0, the promoter is nearly fully repressed, but with increasing *A*, the mean population 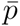 of free TetR dimers is reduced, weakening the repression and boosting the mean mRNA population 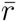. Changing *A* allows us to explore a wide range of control network behavior. Note that since 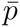 depends on *B* only through the the product *κ*_0_*B*, and the value of this product is fixed at a constant value from the experimental fit (Table 1), 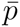 is independent of *B*. On the other hand, 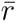, which is proportional to *κ*_0_, is inversely proportional to *B*.

**Table 1:**
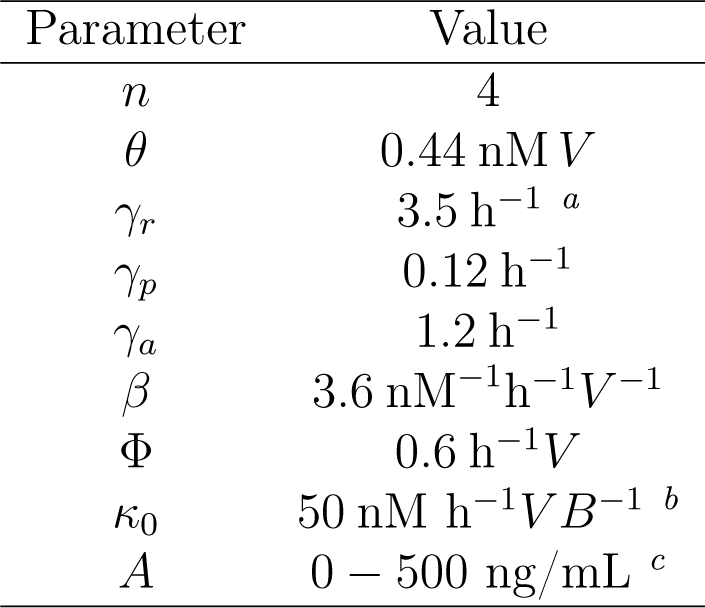
**Parameter values for the dynamical model of the yeast synthetic gene circuit (Eq. (10)). The cell volume *V* is assumed fixed. Unless otherwise noted, all values are taken from the experimental fit of Ref. 24**. ^a^Ref. 31 ^b^The burst ratio *B* = *κ*_*p*_ /*γ*_*r*_. Though not independently determined by the experimental fit, we assume that *B* is in the range *B* = 2 - 10.^30^ ^c^For external ATc concentration *A*, 1 ng/mL corr responds to 2.25 nM.

Linearizing Eq. (10) around 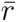 and 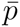, we extract the following frequency-domain response functions:

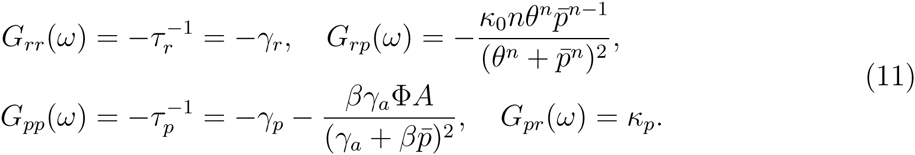

All the functions are constants in the frequency domain. Here *τ*_*r*_ and *τ*_*p*_ are effective decay times for the mRNA and proteins, respectively. The value of *τ*_*r*_ is fixed, and sets the intrinsic time scale of mRNA fluctuations, but *τ*_*p*_ and *G*_*rp*_ depend on 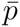, which is a function of the external ATc concentration *A*. In fact, association with intracellular ATc, described by the second term in the *G*_*pp*_ expression above, is the dominant form of decay for the free TetR dimers. Fig. 3(d) plots the effective decay constants 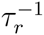 and 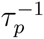 as a function of *A*. Except for *A* ≲ 8 ng/mL we are in the regime where 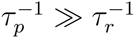, which is relevant in simplifying the optimality condition for *G*_*rp*_*(ω*) discussed below.

The optimal filter calculation for the TetR gene circuit depends on the linear response functions of Eq. (11). We obtain the following power spectra for the signal and noise in the absence of extrinsic noise:

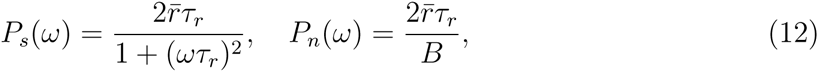

where the burst ratio *B* ≡ *κ*_*p*_*τ*_*r*_ is the mean number of proteins synthesized per mRNA during the lifetime *τ*_*r*_. The problem is to evaluate Eq. (9) for *H*_opt_*(ω*). The sum of signal plus noise, *y(ω) = s(ω) + n(ω*), has a power spectrum *P*_*y*_*(ω*) = *P*_*s*_*(ω*) + *P*_*n*_*(ω*), which we canewrite as follows:

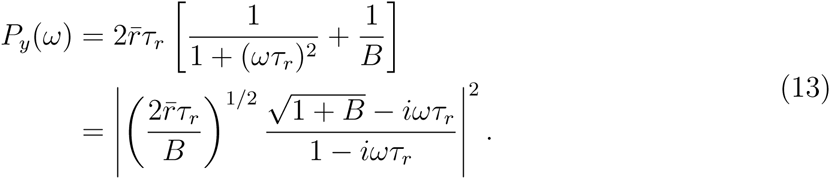

The expression within the absolute value brackets is zero only at 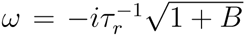, and has a simple pole at 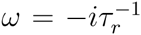. Since all the zeros and poles are in the lower complex ω half-plane, it satisfies the criterion for the causal term in the factorization 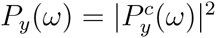. Thus:

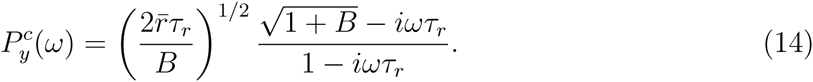

The other causal term in Eq. (9) involves the additive decomposition 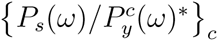. This is calculated by looking at the partial fraction expansion of 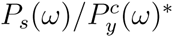:

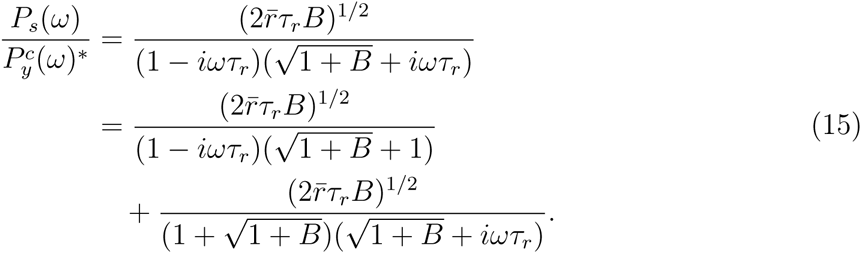

Of the two terms in the partial fraction expansion, only the first has poles solely in the lower complex ω half-plane. Hence, it is the only one that contributes to 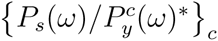:

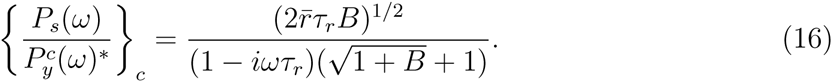

Inserting Eqs. (14) and (15) into Eq. (9), we finally find that the optimal filter is:

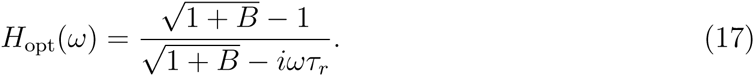

Transforming *H*_opt_*(ω*) into the time domain, we find

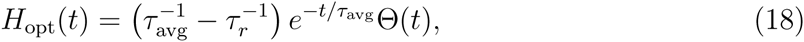

where 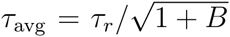, and Θ(*t*) is a unit step function ensuring that the filter operates only on the past history of its input. For B≫1 the prefactor in Eq. (18) is 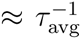, and *H*_opt_*(t*) has a straightforward interpretation: it approximately acts as a moving average of the corrupted signal *y(t) = s(t*)+*n(t*) over a time scale *τ*_avg_. In order to get the best estimate 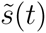, the averaging interval *τ*_avg_ can neither be too long, since it would blur out the features of the signal *s(t*) (which vary on the time scale *τ*_*r*_), nor too short, since it would be ineffective at smoothing out the noise distortion *n(t*). Hence, there must exist an optimum *τ*_avg_, which is naturally proportional to τ_*r*_, the main time scale for the mRNA.

In Fig. 3(b), we show how the noise filter properties of the system vary with *A* for a burst ratio of *B* = 10. The filter function *H(t*) (solid red curve) differs substantially from *H*_opt_*(t*) (dotteded curve) for large and small *A*, but approaches the optimal form near *A* = 54 ng/mL. Consequently, at this value of *A* we get the closest correspondence between the plotted sample trajectories of signal *s(t*) (cyan curve) and estimate 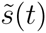 (blue curve). Similarly, the equilibrium probability distribution of the output, *P(δr*), shown to the right of the trajectories, exhibits the smallest Fano factor 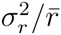. The latter is a measure of noise magnitude, and has a reference value of unity if mRNA production was a pure Poisson process, as would be the case without feedback. Optimality is realized in the intermediate *A* regime of partial repression, where the *R* to *P* responsiveness, as measured by |*G*_*rp*_|, is large. Effective noise suppression requires that *R* be sensitive to changes in *P*, so that information about *R* fluctuations can be transmitted through the negative feedback loop.

In order to understand the optimality condition for *H(t*) in more detail, let us look at the explicit expression for *H(t*) in the TetR system, given by the inverse Fourier transform of Eq. (6) with the response functions of Eq. (11):

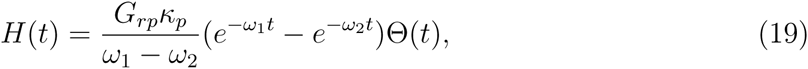

where *ω*_1_, *ω*_2_ are the two *ω* roots of the denominator in Eq. (6). Assuming *τ*_*p*_≪ *τ*_*r*_ (which holds good except for small values *A* ≲ 8 ng/mL, as seen in Fig. 3(d)), we can directly show the approach of *H(t*) to optimality at a specific intermediate value of *G*_*rp*_. When *G*_*rp*_ equals 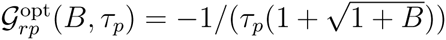, the roots 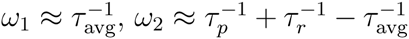, up to corrections of order 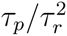. In this case, Eq. (19) becomes

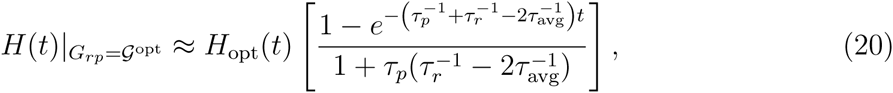

where the factor in the brackets on the right equals 1 in the limit *τ*_*p*_ → 0 for all *t* > 0. Up to this correction factor, we thus expect the system to behave optimally at *A* = *A*_opt_, defined by the condition 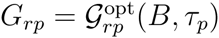, so long as *A*_opt_ is large enough to satisfy *τ*_*p*_ ≪ *τ*_*r*_. Fig. 3(e) shows *G*_*rp*_ and 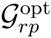 curves for *B* = 2, 5, 10, with dots marking the intersection points that define *A*_opt_ for each *B*. As explained above, |*G*_*rp*_| is small at small and large *A*, and reaches a maximum in between. At fixed *B*, 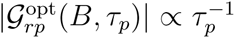, so it increases monotonically with *A*, as larger concentrations of the inducer increase the effective decay rate of free proteins. Thus, for each *B* there is a single intersection point *A*_opt_ at an intermediate concentration of the inducer.

Fig. 3(f) shows the Fano factor 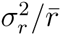 versus *A* for various *B*. As the control network approxim rates optimality at *A*_opt_ for each *B*, the Fano factor nears its minimum, close to the theoretical limit marked by the horizontal dashed lines. This limit is the minimal possible 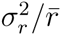, calculated from Eq. (8) using *H*_opt_*(t*) from Eq. (18):

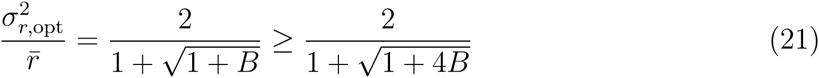

A few comments concerning the above equation are in order. (1) The result on the far right-hand side is the rigorous lower bound derived by LVP.^19^ In their case, the feedback mechanism through the rate function *κ*_*r*_*(t*) could be any causal functional of *p(t*), linear or nonlinear. The Fano factor of the optimal linear filter differs in form only by the coefficient of *B*, and is always within a factor of 2 of the lower bound for any value of *B*. (2) For Gaussian-distributed signal *s(t*) and noise *n(t*) time series, the linear filter is optimal among all possible filters.^28^ If the system fluctuates around a single stable state, and the copy numbers of the species are large enough that their Poisson distributions converge to Gaussians (mean populations ≳ 10), the signal and noise are usually approximately Gaussian. This is a wide class of systems where the rigorous lower bound (the last term in Eq. 21) can never be achieved. In other words, here the WK filter yields the most efficient feedback mechanism. Although, as pointed out by LVP, nonlinearity could lead to additional noise reduction, the benefits are likely to be restricted to those systems where the signal and/or noise are substantially non-Gaussian. However, since the form of the optimal control network has not been found in the general nonlinear case, it remains an interesting open question whether the LVP bound can actually be reached even within this category of systems. We willeturn to this issue in the next section. (3) The parameter *B* is the key determinant of noise reduction. For B ≪ 1, there are not enough signaling events to control the mRNA fluctuations, and as *B* → 0 we approach 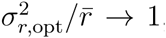, the no-feedback Poissonesult. In the limit *B* ≫ 1 signaling is effective, and the Fano factor decreases with *B* as 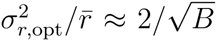. For large enough *B* we approach perfect control, but at extreme expense: the standard deviation of the mRNA fluctuations *σ*_*r*,opt_ ∝ *B^−1/4^*, the same scaling derived by LVP.

## WK theory constrains the performance of a broad class of nonlinear, discrete regulatory networks

The results in Fig. 3 rely on a linearized, continuum approach to the TetR dynamical system. To assess if the conclusions based on the WK optimal filter hold if these approximations are relaxed, we first performed kinetic Monte Carlo simulations of the full nonlinear system (Eq. (10)) using the Gillespie algorithm.^32^ We chose a cell volume of *V = V*_0_ = 60 fL, within the observed range for yeast,^33^ which corresponds to the mean populations 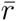 and 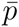 shown in Fig. 4(a) as a function of *A*. (For example, at *A = A*_opt_ = 62.7 ng/mL when *B* = 5, 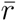 ≈ 84 and 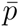 ≈ 11. In addition to the nonlinearity, the discrete nature of the populations in the simulation might play a role at these low copy numbers.) The numerical results for the Fano factor 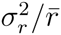 are plotted in Fig. 4(b) at *B* = 2, 5, 10, for *V = V*_0_ (circles) and also for comparison at a larger volume *V* = 10*V*_0_ (squares). The blue curves show the linear theory results, and the dashed lines are the optimality predictions for 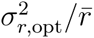. Although nonlinearity and discreteness effects do change the results, the linear theory gives a reasonable approximation, and the minimum is still near *A*_opt_. The feedback mechanism is nonlinear in the simulations, but it does not do better than the linear predictions for 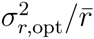 for the parameters used to describe the experimental results. Though the intrinsic population noise is Poisson-distributed in the simulations, the Poisson distribution is very close to Gaussian, even for copy numbers as low as ~ 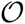. Since the linear filter is the true optimum for a Gaussian-distributed signal and noise,^28^ we do not expect improvements in noise suppression by employing a nonlinear version. In the opposite limit of large copy numbers, *V* → ∞, the continuum approximation should be valid, and population fluctuations increasingly negligible relative to the mean. Thus, the linear theory should directly apply in this limit, and indeed we see that for *V* = 10*V*_0_ the discrepancies between numerical and theory results are substantially reduced (Fig. 4(b)). It is worth emphasizing, that even at the realistically small cell volume *V*_0_, the linear theory retains much of its predictive power. More generally, the conditions for WK optimality do not have to be perfectly satisfied in order for the filter to perform close to maximum efficiency There is an inherent adaptability and robustness in near-optimal networks, as reflected in the broad minima of 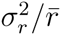 as a function of *A* (Fig. 4(b)).

**Figure 4:**
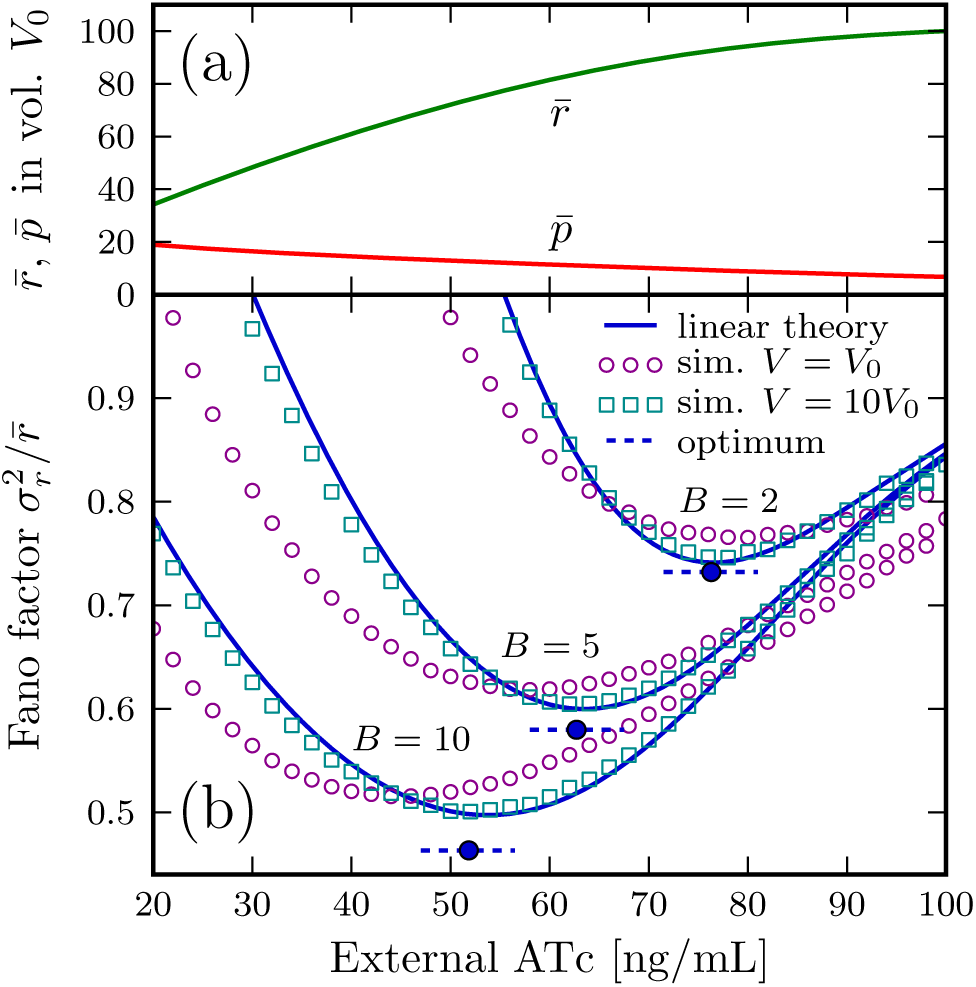
Results of simulation and theory for the yeast synthetic gene circuit,^24^ as a function of extracellular ATc concentration *A*. (a) Mean populations of free TetR mRNA 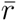 and TetR dimer 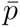, assuming a cell volume *V*_0_ = 60 fL. (b) The Fano factor 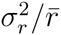 for burst factor *B =* 2,5,10, as predicted by the linear filter theory (solid lines), versus stochastic numerical simulations at two different volumes, *V = V*_0_ (circles) and *V* = 10*V*_0_ (squares). The WK filter theory predicts the minimal Fano factor 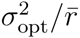 given by Eq. (21) (horizontal dashed lines). The system can be tuned to approach optimality near a particular *A*_opt_ obtained by the condition 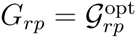 (filled circles).

The semi-quantitative agreement between the linearized theory and the simulation results displayed in Fig. 4 still leaves open the possibility that some type of nonlinear, discrete filter, not described by the experimentally fitted parameters of the TetR gene network, could perform better than the WK optimum at sufficiently small volumes. Fig. 5 plots both the WK value for the Fano factor (solid curve) and the rigorous lower bound of LVP (dashed curve) as a function of *B* (Eq. (21)). The above question can be posed as follows: is it possible to achieve a Fano factor that falls between the two curves by taking advantage of nonlinearity and discreteness? Ideally, one should do an optimization over all possible nonlinear regulatory functions that could describe feedback between the TetR protein and mRNA. In full generality, such an optimization appears intractable, but one can tackle a limited version of the nonlinear optimization. We will confine ourselves to Hill-like regulatory functions, which describe the experimental behavior of many cellular systems,^34^ and explore whether it is possible to find any scenario where this type of nonlinear feedback outperforms the linear WK optimum. We consider the following generalized TetR feedback loop:

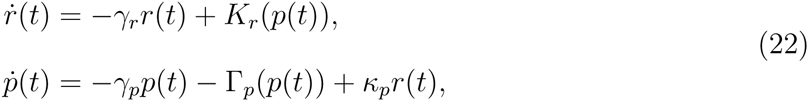

with two Hill-like regulatory functions,

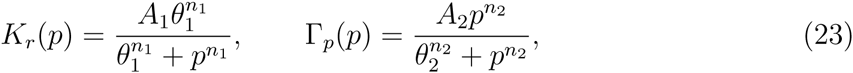

involving arbitrary non-negative parameters *A*_*i*_, *n*_*i*_, *θ*_*i*_, *i* = 1, 2. The original TetR system (Eq. (10)) is a special case of the equations above with:

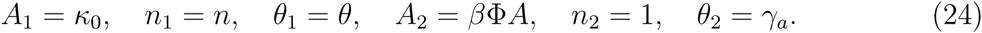

**Figure 5:**
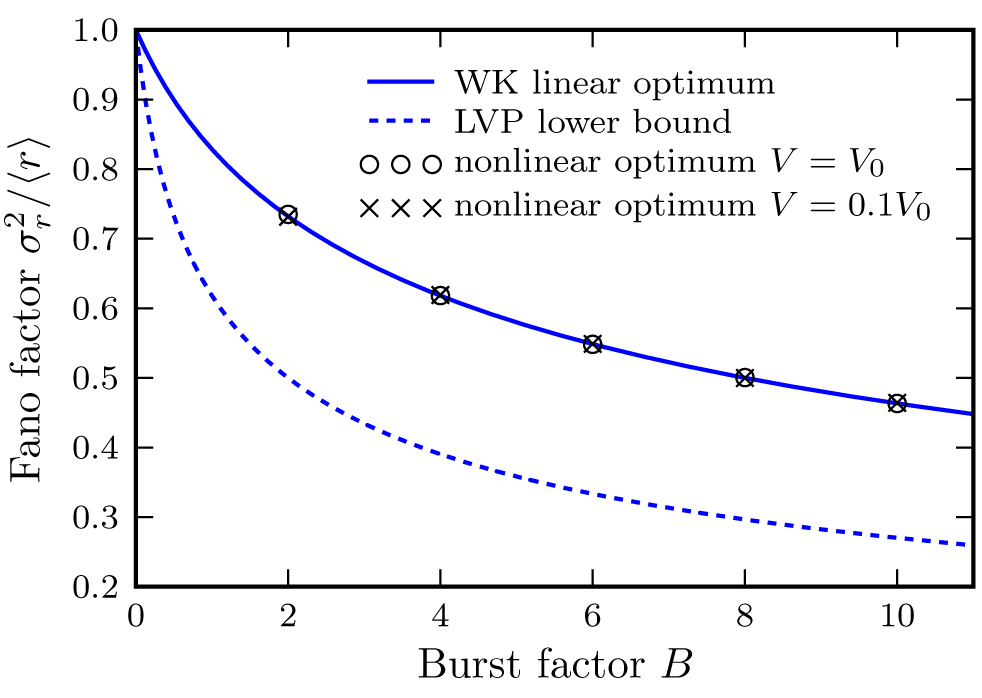
The Fano factor 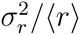 as a function of burst ratio *B*. The solid curve is the optimal result predicted by the WK linear theory, and the dashed curve is the rigorous lower bound derived by LVP.^19^ Symbols show numerical optimization for the generalized nonlinear TetR feedback system (Eq. (22)) at two volumes, *V = V*_0_ and *V* = 0.1*V*_0_.

The production function *K*_*r*_*(p*) is a monotonically decreasing function of *p*, as is expected for negative feedback, while *Γ*_*p*_*(p*) is monotonically increasing, a generalization of some regulatory network which effectively removes the TetR protein from the feedback loop (the role played by ATc binding in the experimental system). With these monotonicity constraints, there is always only one steady-state solution 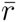 and 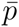 to Eq. (22).

The optimization consists of searching for *K*_*r*_*(p*) and *Γ*_*p*_*(p*) that minimize the Fano factor 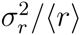. The following quantities are fixed during the search: the degradation rates *γ*_*r*_, *γ*_*p*_, the *P* production rate *κ*_*p*_ (or equivalently the burst ratio *B = κ*_*p*_/*γ*_*r*_), and the steady state values 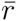, 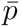. Note that in the general nonlinear case, the steady state values do not necessarily coincide with the mean values 〈*r*〉, 〈*p*〉, since the equilibrium distributions are generally asymmetric with respect to the steady state. Fixing 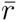 and 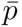 during the optimization is one way to set an overall copy number scale, to investigate the role of discreteness. It turns out that the optimization results described below end up being independent of 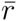 and 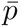. In terms of the Hill function parameters, fixing 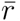 and 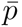 means setting *A*_1_ and *A*_2_ to the following values,

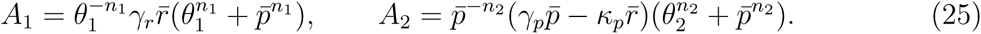

Thus the goal of optimization is to minimize 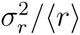 over the four remaining free parameters: *n*_1_, *θ*_1_, *n*_2_, *θ*_2_.

In order to carry out this minimization, one needs an efficient procedure to calculate 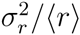 from Eq. (22), keeping both the full nonlinearity of the dynamical system, and the discreteness of the *r(t*) and *p(t*) populations. The system can always be simulated through the Gillespie algorithm,^32^ and accurate estimates of 〈r〉 and 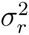 determined from sufficiently long trajectories. However this approach is too slow for searching over the four-dimensional parameter space, since each distinct set of parameters would require a separate long simulationun. An equivalent, faster alternative is to directly solve the system’s master equation for the steady state probability distribution, which then yields 〈r〉 and 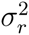. The joint probability distribution *P*_*r,p*_*(t*) of finding mRNAs and *p* proteins at time *t* is governed by the master equation,

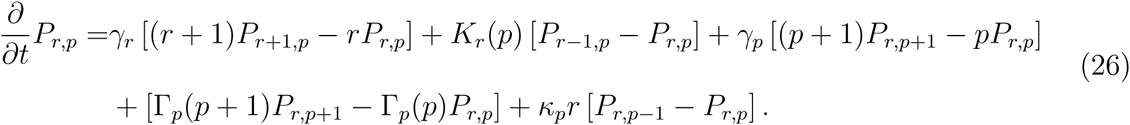

The steady state distribution 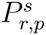, is the solution obtained by setting to zero the right-hand side of the above equation, which we denote *R* _*r,p*_:

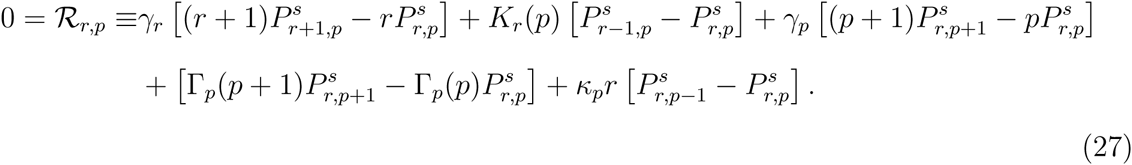

The result is linear in the components 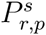 for various *r* and *p*, and thus the set {*R*_*rp*_ = 0} for = 0,1,… and *p =* 0, 1,…, constitutes a linear system of equations for 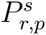. The master equation can be solved by spectral methods, which are generally more efficient than brute force Gillespie simulations.^35^ However we use a different approach, described below, to solve Eq. (27), which is sufficiently fast for our numerical optimization purposes. Since *r* and *p* can take on any integer values between 0 and ∞, we truncate the system to focus only on the non-negligible 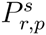, in other words *(r,p*) within several standard deviations of the mean (〈*r*〉,〈*p*〉). Specifically, we keep only those equations *R*_*rp*_ = 0 which involve *r*_min_ ≤ *r* ≤ *r*_max_ and *p*_min_ ≤ *p* ≤ *p*_max_. The largest truncation range required for accurate results was *r*_max_ - *r*_min_ = 100 and *p*_max_ - *p*_min_ = 50. All 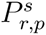 outside the range which appear in the truncated system of equations are set to a positive constant ∊ > 0. (The precise value of ∊ is unimportant since the distribution is subsequently normalized, and the truncation range is chosen large enough so that the boundary condition does not significantly affect the outcome.) The resulting finite linear system, which is sparse, can be efficiently solved using an unsymmetric-pattern multifrontal algorithm.^36^ Knowing 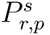, we then directly calculate the moments of the distribution to find 〈*r*〉 and 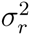. The numerical accuracy of the procedure is verified by comparison to Gillespie simulation results.

In order to set a starting point for each round of nonlinear optimization, we use the following initialization procedure: we take the original TetR system at a given volume *V* and burst ratio *B* (fixing the Hill function parameters according to Eq. (24)) and find the ATc concentration *A*_min_ where 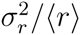 is smallest, evaluating the Fano factor using the linear solver described above. The 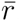 and 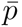 at this concentration are then chosen to be fixed constants for the nonlinear optimization, where we vary the parameters *n*_1_, *θ*_1_, *n*_2_, *θ*_2_ from the initial values given by Eq. (24) to minimize 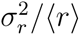. The minimization is carried out using Brent’s principal axis method,^37^ which is feasible due to the fast evaluation of 〈*r*〉 and 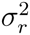 at each different parameter set through the linear solver.

Fig. 6 shows results of a typical minimization run, where the initial system is at volume *V* = *V*_0_ with *B =* 10, with a corresponding *A*_min_ = 50 ng/mL. The dashed lines in Fig. 6(a) and (b) show the Hill functions *K*_*r*_*(p*) and *Γ*_*p*_*(p*) of the original TetR system at these parameter values, and the heat map in Fig. 6(c) represents the associated steady-state probability distribution 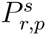. The dashed lines superimposed on the heat map are the loci of solutions to *ṙ(t*) = 0 and *ṗ(t*) = 0 (the right-hand sides of Eq. (22) set to zero), which intersect at the steady state 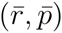. The Fano factor for this distribution, which represents the best the TetR system can perform given the experimentally fitted parameters, is 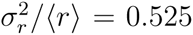. This is above the linear WK optimum for 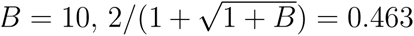, and significantly larger than the rigorous LVP lower bound of 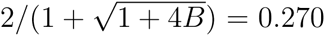. Once we relax the experimental constraints, and carry out the numerical minimization, the Fano factor decreases. The solid lines in Fig. 6(a) and (b) show *K*_*r*_*(p*) and *Γ*_*p*_*(p*) after several steps of the minimization algorithm, and Fig. 6(d) shows the corresponding 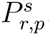. The Hill functions have become very steep steps around 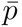, while the average of the distribution 〈*r*〉 has been pushed above 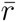. The probabilities 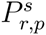 for *p* < *p*_0_ become negligible, where 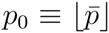 is the largest integer value below 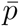. For *p* > *p*_0_, 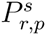 rapidly decay to zero. The Fano factor, 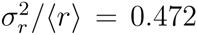, approaches closer to the linear WK optimum, but is still above it. If we allow the minimization to proceed, these trends continue: at each iteration the Hill functions get steeper, 〈*r*〉 increases, 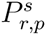 for *p* < *p*_0_ tends to zero, and 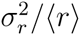 approaches arbitrarily close to the linear WK optimum from above.

**Figure 6:**
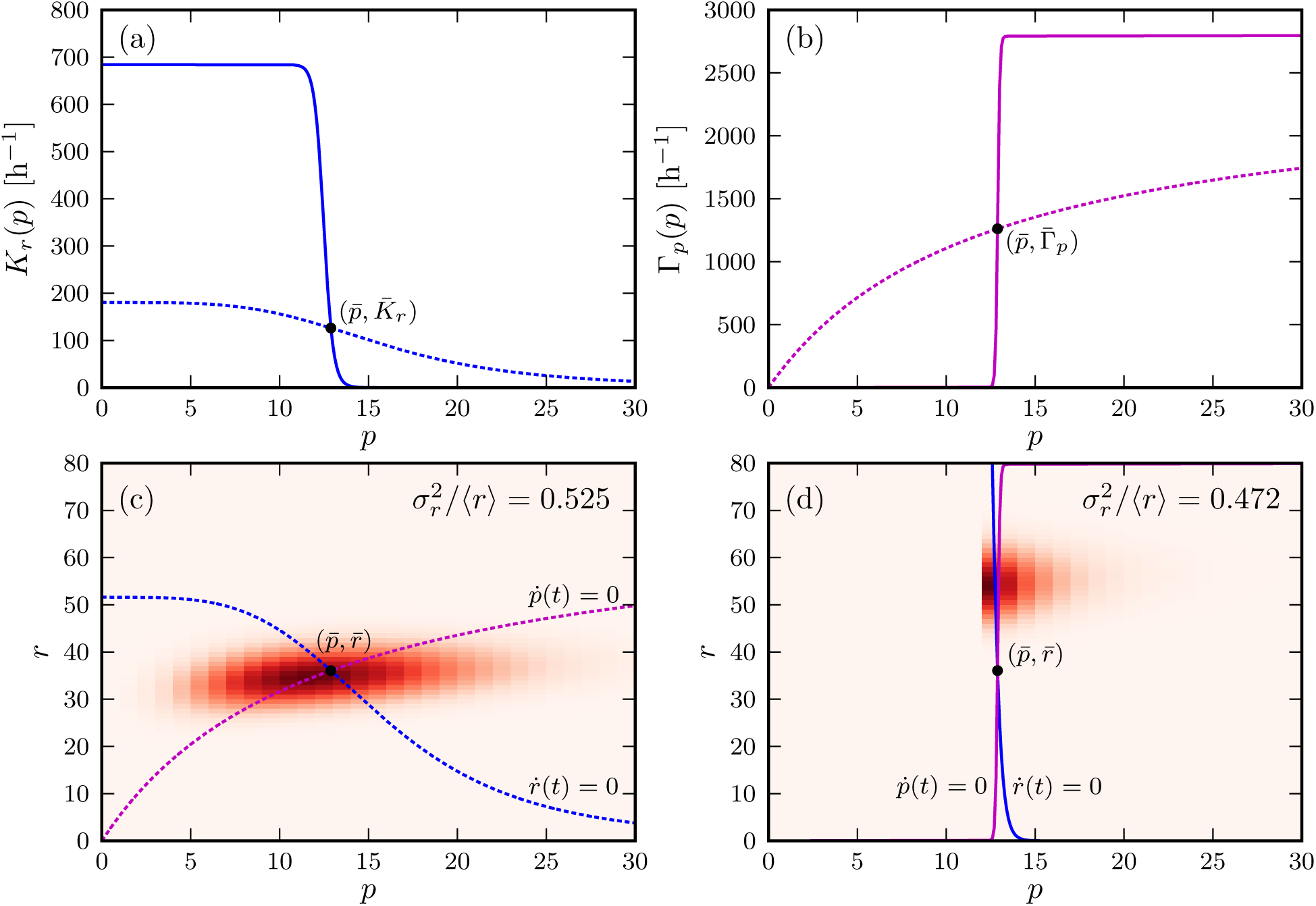
Results for numerical optimization of the generalized nonlinear TetR feedback system of Eq. (22), with starting parameters *B =* 10 and *V* = *V*_0_. (a) The mRNA production regulation function *K*_*r*_*(p*) in its initial form before optimization (dashed curve), and after several steps of the minimization algorithm (solid curve). (b) Similar to (a), but showing the protein degradation function *Γ*_*p*_*(p*). (c) Heat map of the steady-state probability distribution 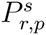 before optimization, corresponding to regulation governed by the dashed curves in the top panels. The nullclines *ṙ(t*) = 0 and *ṗ(t*) = 0 are superimposed. (d) Similar to (c), but after several steps of the minimization algorithm, corresponding to regulation governed by the solid curves in the top panels.

In fact, the same behavior is seen irrespective of the volume *V* and burst ratio *B* used to define the initial point of the optimization. Fig. 5 shows the results of nonlinear optimization for *B* = 2 - 10 at two volumes, *V* = *V*_0_ and *V* = 0.1*V*_0_. Even for the smallest volume, the nonlinear optimization results can get arbitrarily close to the WK optimum, but never do better. No generalized nonlinear system based on Hill function regulation brings us close to the theoretically possible LVP lower bound. This overall conclusion holds even when we change the functional form for the generalized feedback. We tried two alternatives: (i) using sigmoidal (logistic) functions instead of Hill functions; (ii) expanding *K*_*r*_*(p*) and *Γ*_*p*_*(p*) in a Taylor series around 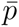, truncating after the third order term, and minimizing with respect to the Taylor coefficients. In both cases numerical minimization of the Fano factor led to similar step-like behavior for *K*_*r*_*(p*) and *Γ*_*p*_*(p*), and the Fano factor tended to WK optimum from above.

From the 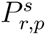 distribution in Fig. 6(d) we see that the step-function limit leads to a system which is highly nonlinear along the *p* axis: in fact the gene network spends most of its time at *p* = *p*_0_, just below the sudden change in regulation due to the steep Hill functions, and *p* > *p*_0_ just above the sudden regulatory change. The feedback on the TetR mRNA population is mediated by *p* fluctuations between the two regimes, resulting in threshold-like regulatory behavior. Remarkably, despite this discrete, nonlinear character, the network can still approach the efficiency of an optimal WK linear filter. To gain a deeper understanding of how the step-like regulation can match WK optimality, we used the numerical optimization results described above to posit a limiting form of the nonlinear gene network that can be solved analytically (details in Appendix C). The analytic results explicitly show that we can asymptotically approach the WK optimum behavior from above, even in systems where the protein copy numbers are very small. Thus at least for a two-component TetR-like system regulated by biologically-realistic Hill functions, the constraint derived from the WK theory has a broader validity than one would guess from the underlying continuum, linear assumptions. It thus becomes an interesting and a non-trivial problem, left for future studies, to find an example of a gene network where the rigorous lower bound of LVP could be directly achieved.

## Realizing optimality under the influence of extrinsic noise

Extrinsic noise is ubiquitous and hence must also be considered in any effective description of the control network. Inevitably, certain cellular components are not explicitly included in such a description, which in our case study could include RNA polymerase, ribosomes, and transcription factors that bind to the same promoter. Each of these components have their own stochastic characteristics and may contribute noise to a smaller or greater extent. Particularly for eukaryotes like yeast, the extrinsic noise contribution may be significantly larger than the intrinsic component.^38^,^39^ We adopt a simple model for the extrinsic noise based on earlier approaches,^14^,^16^ which assume that it is band-limited at a low frequency 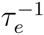, where *τ*_*e*_ is on the order of the cell growth time scale. The justification is that higher frequency contributions to the extrinsic noise are filtered out by the gene circuits associated with its sources. This idea is consistent with the experimental observation of extrinsic noise in protein production in *E. coli*, which found long autocorrelation times for the extrinsic noise on the order of the cell cycle period.^40^

For the TetR system, our theory is extended to the extrinsic noise case in Appendix D, with the results illustrated in Fig. 7. The outcome is that a given TetR gene circuit, tuned appropriately such that *A = A*_opt_, can act as a WK filter for an entire family of extrinsic noise scenarios. A single set of parameters can approximately represent the optimal solution for a variety of extrinsic inputs. This makes the WK concept a versatile design tool for noise suppression in biological systems: the same control network can act with maximum efficiency in a variety of different contexts. It is possible that the requirement of adaptability to a wide range of conditions has resulted in the evolution of control networks acting as WK filters. It remains to be seen whether nature has exploited this feature *in vivo*.

**Figure 7:**
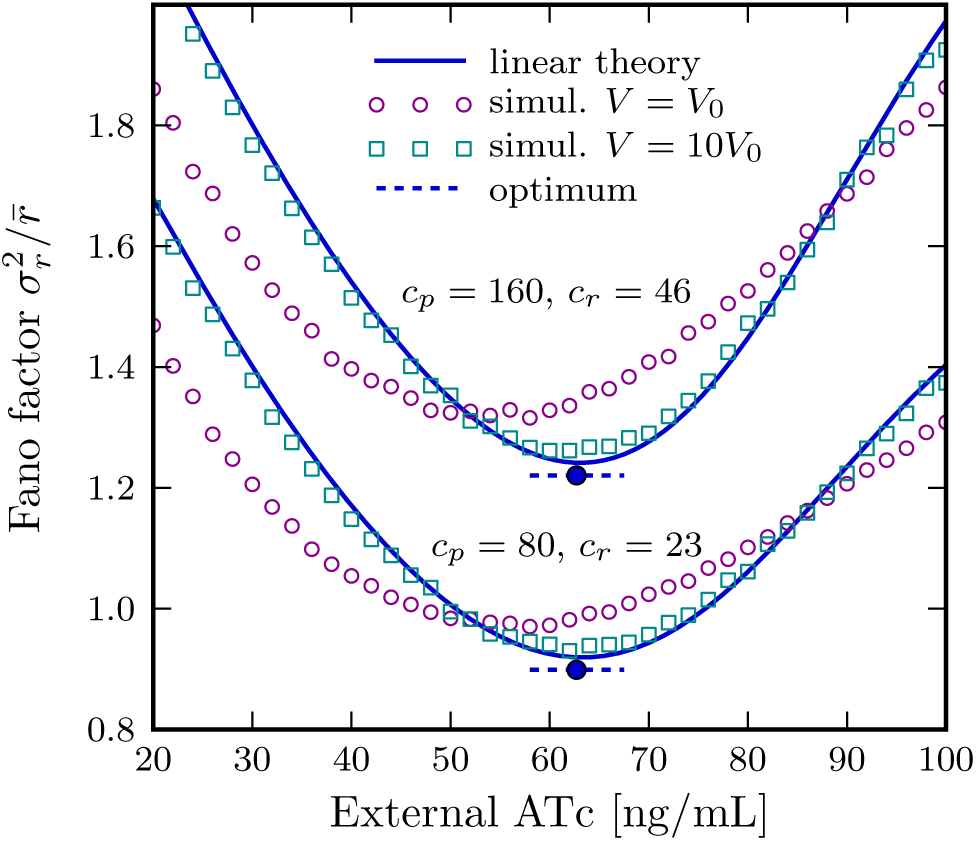
Comparison of simulation and theory results based on the dynamical model (Eq. (10)) of the yeast synthetic gene circuit,^24^ in the presence of extrinsic noise given by Eq. (73). All quantities are plotted as a function of extracellular ATc concentration *A* for the burst ratio *B* = 5. Each set of curves shows the Fano factor 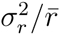, as predicted by the linear filter theory (solid lines), versus stochastic numerical simulations at two different volumes, *V* = *V*_0_ = 60 fL (circles) and *V* = 10*V*_0_ (squares). The two sets correspond to noise magnitudes *c*_*p*_ = 80, *c*_*r*_ = 23 and *c*_*p*_ = 160, *c*_*r*_ = 46. In both cases *c*_*r*_ and *c*_*p*_ are related through the condition in Eq. (83), and the minimal Fano factor predicted by WK filter theory (horizontal dashed lines) is modified as shown in Eq. (84). The system can be tuned to approach optimality near a particular *A*_opt_ obtained by the condition 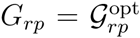(filled circles).

## Conclusion

The TetR feedback loop is a concrete example of how a WK filter can be implemented in a gene network driven by a complex set of biochemical reaction rates, but the overall approach outlined here has far reaching implications, thus highlighting the appeal of engineering paradigms in biology.^41^ With the entire network complexity encoded in a handful of response functions, we can derive fundamental limits and design principles governing biological regulation. The key step is to map the linear response picture onto a signal estimation problem, whose solution is given by WK theory. This idea allows us to predict the dynamic properties of the feedback pathway required to optimally filter noise in a broad class of negative feedback circuits. As already demonstrated in earlier works,^22^,^23^ the mapping, and the potential utility of the WK approach, is not unique to the negative feedback loop. Another important byproduct of the theory is that the behavior of gene circuits away from optimality can also be predicted. In this sense, our practical approach goes beyond just obtaining rigorous bounds, and allows us to characterize how close or far gene networks are from optimality for biologically relevant parameters.

We have derived response functions by linearizing a minimal model extracted from experimental observations, but it is also possible to directly apply small perturbations to a system, and measure the resulting time-dependent changes in populations of species. Recently, the yeast hyperosmolar signaling pathway has been probed by perturbations in the form of salt shocks.^42^^-^^44^ Despite the underlying complex nonlinear network, the details of which are not completely characterized, a linear response description quantitatively captures the frequency-dependent behavior of the pathway over a wide range of inputs. *E. Coli* chemotaxis signaling also exhibits a linear regime,^45^ where the fluctuation-dissipation relationship between the system’s unperturbed behavior and its reaction to external stimuli has been explicitly verified.

Linear response functions can thus become a fundamental tool in analyzing biochemical circuits, analogous to their established role in control engineering and signal processing. More extensive experimental measurements will be critical in this effort, in order to ascertain how varied the response relationships between regulatory components are in nature. Once we understand the essential dynamical building blocks out of which complex biological function is realized, we can map out the hidden constraints that control the behavior of living systems.

## Acknowledgement

This work was done while the authors were in the Institute for Physical Sciences and Technology in the University of Maryland, College Park. We are grateful to C. Güven, G. Reddy, Z. Zhang, and P. Zhuravlev for useful discussions. This work was supported by a grant from the National Science Foundation (CHE 13-61946).

## Appendix A: Derivation of the optimal WK filter

In this section we derive Eqs. (8) and (9) in the main text. They describe the output variance 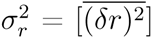 and the linear filter *H*_opt_(*ω*) that minimizes 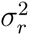, which are the main quantities in the Wiener-Kolmogorov theory.

**Output variance** 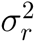 **in terms of signal and noise power spectra *P*_*s*_(*ω*) and *P*_*n*_(*ω*)**

From Eq. (4), which defines the signal *s(ω*) and estimate 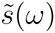 in the frequency domain, the Fourier transformed output *δr(ω*) for any *H(ω*) can be rewritten as,

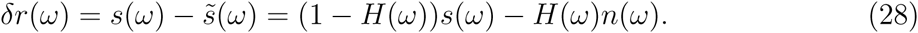

In the time domain, *s(ω*) = -*n*_*r*_(*ω*)/(*G*_*rr*_(*ω*)+*iω*), is a convolution of the noise function *n*_*r*_*(t*), and *n(ω*) = *n*_*p*_(*ω*)/*G*_*pr*_(*ω*) is a convolution of *n*_*p*_*(t*). So long as the noise functions *n*_*r*_*(t*) and *n*_*p*_*(t*) are uncorrelated, *s(t*) and *n(t*) are also uncorrelated, so the frequency domain average 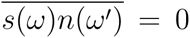. (The theory can also be generalized to correlated noise sources, but for simplicity we consider only the uncorrelated case.) As a result, the correlation 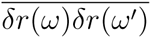, related to the output power spectrum *P*_*δr*_(*ω*), can be written in terms of *P*_*s*_(*ω*) and *P*_*n*_(*ω*), the individual power spectra of the signal and noise:

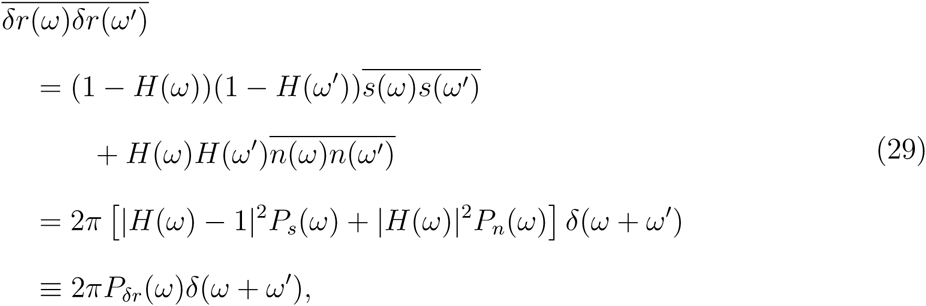

In the above equation we have used the definition of the power spectrum, i.e. 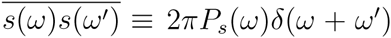, and the relation *H(-ω*) = *H*(ω*) since *H(ω*) is the Fourier transform of a real function *H(t*). The power spectrum *P*_*δr*_(*ω*) is the Fourier transform of the time autocorrelation function 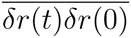:

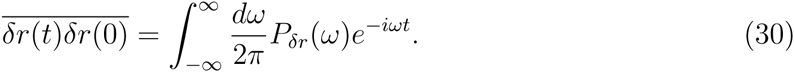

At *t* = 0, the autocorrelation function gives us the variance 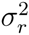:

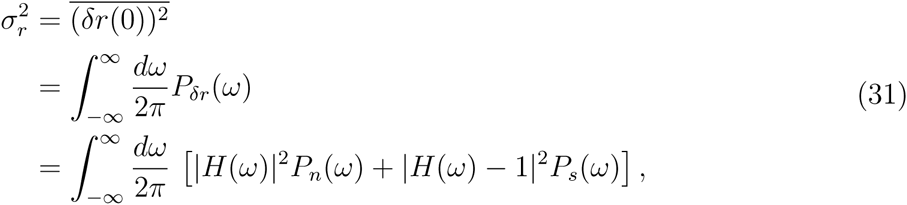

which is Eq. (8) in the main text.

**Minimizing** 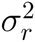 **over all causal *H(ω*) yields the optimal WK filter *H*_opt_(*ω*)**

The convolution of the filter function *H(t*) on the corrupted signal *s(t*) + *n(t*) must satisfy causality. The filter can only operate on the past history of *s(t*) + *n(t*), so *H(t) =* 0 for *t<*0. In the frequency domain, enforcing causality restricts *H(ω*) to have certain general properties as a function of complex *ω*:^29^ it can have no poles or zeros in the upper half-plane Im*ω*>0. Equivalently, the real and imaginary parts of *H(ω*) evaluated at real *ω* must satisfy the well-known Kramers-Kronig relation:

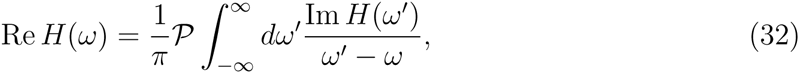

where *P* is the Cauchy principal value of the integral. The goal of WK optimization is to minimize 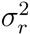 in Eq. (31) over all possible causal functions *H(ω*), given the power spectra *P*_*s*_*(ω*) and *P*_*n*_*(ω*).

Assume such an optimum *H*_opt_*(ω*) exists, with the corresponding minimal variance 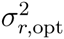. Let us add a small perturbation, *H(ω*) = *H*_opt_(*ω*) + *δH(ω*), where *δH(ω*) is also a causal function of complex *ω*. From Eq. (31), the resulting variance change 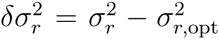, to lowest order in *δH(ω*), is:

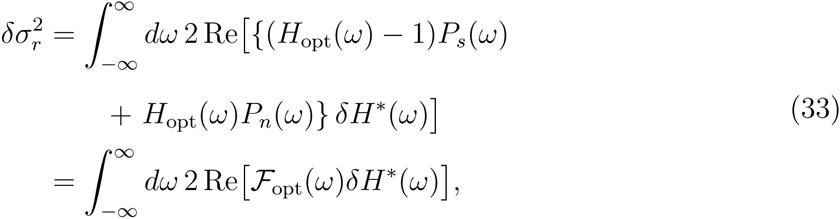

where

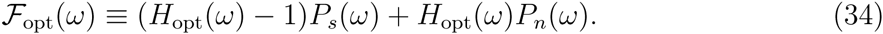

For *H*_opt_*(ω*) to be the WK optimum, 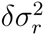 in Eq. (33) must be zero for any causal perturbation *δH(ω*).

Out of all possible causal perturbations, we will focus on one with the specific form:

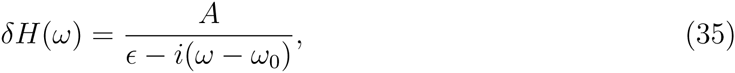

where Im*ω*_0_ = 0 and *A, ∊*>0. It has no zeros, and the only pole, *ω = ω*_0_ - *i∊*, is in the lower half-plane, so *δH(ω*) is causal. We will be interested in the limit as this pole approaches the real axis, *∊* → 0^+^, where the real and imaginary parts of *δH(ω*) are,

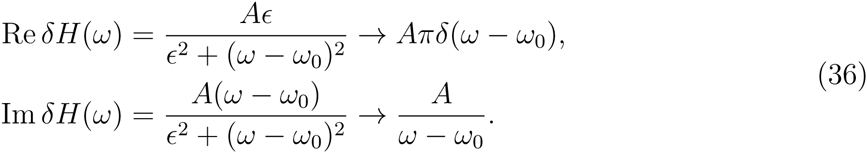

Substituting these into Eq. (33) for 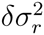, we find that the optimality condition 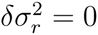 implies the following relation between the real and imaginary parts of *F*_opt_(*ω*):

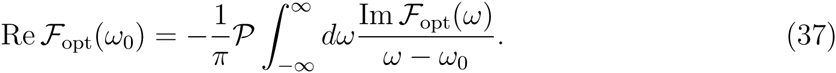

This has the same form as the Kramers-Kronig relation in Eq. (32), with the important difference of a minus sign in front. Consequently, *F*_opt_(*ω*) must be *anticausal*, which we define as a function with no poles or zeros in the lower complex ω half-plane.

In order to use this result to derive a solution for *H*_opt_(*ω*), we define two types of decompositions, described briefly in the main text. In practice, all the frequency domain power spectral density and filter functions we work with in the linear response formalism are meromorphic functions over the complex *ω* plane. Any meromorphic function *F(ω*) can be written as a partial fraction expansion of the form *F(ω*) = Σ_*n, k*_ *c*_*ik*_/*(ω - ω*_*n*_)^*k*^, where {*ω*_*n*_} is the set of poles of *F(ω*), and *c*_*ik*_ are constants. Most generally, the expansion could include a polynomial term, but the functions *F(ω*) we encounter have well-defined inverse Fourier transforms, which require |*F(ω*)| → 0 as |*ω*| → ∞ (decay at least as fast as 1/|*ω*|). Thus, all the terms in the expansion are of the form *c*_*ik*_/(*ω - ω*_*n*_)^*k*^, and we can segregate them according to whether the pole *ω*_*n*_ is in the upper half plane. The causal part {*F(ω*)}_*c*_ is defined as all those terms where *ω*_*n*_ is not in the upper half plane, and the anticausal part {*F(ω*)}_*ac*_ contains the remaining terms in the expansion. The overall function *F(ω*) = {*F(ω*)}_*c*_ + {*F(ω*)}_*ac*_.

The second type of decomposition, an example of Wiener-Hopf factorization,^20^ concerns power spectral density functions like *P*_*y*_(*ω*), which are meromorphic and also real-valued on the real *ω* axis. Let us factor *P*_*y*_(*ω*) as the product of two meromorphic functions, 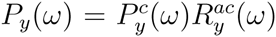. The function 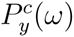 contains all the zeros and poles in *P*_*y*_(*ω*) which are not in the upper half plane. Such a decomposition is always possible, since a meromorphic function can always be written as a ratio of two holomorphic functions. Hence, the numerator and denominator of *P*_*y*_(*ω*) can be decomposed individually into a product of elementary factors by the Weierstrass factorization theorem, with each factor containing a single zero. Because *P*_*y*_(*ω*) is real for real *ω*, so *P*_*y*_*(ω*)^*^ = *P*_*y*_(*ω*) when Im*ω* = 0. Thus, 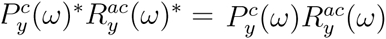. Since 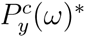 for real *ω* has all its zeros and poles in the upper half plane, we must have 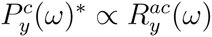, and similarly 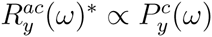. By appropriately absorbing an overall constant into 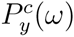, we can factor *P*_*y*_(*ω*) as 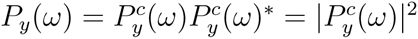.

With these decompositions defined, we return now to the condition in Eq. (37), which shows that *F*_opt_(*ω*) is anticausal. Thus, its causal part in the additive decomposition must be zero, {*F*_opt_(*ω*)}_*c*_ = 0. From the definition of *F*_opt_(*ω*), Eq. (34), it follows that

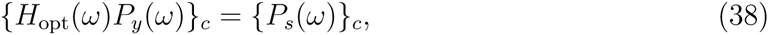

where *P*_*y*_(*ω*) = *P*_*s*_(*ω*) + *P*_*n*_(*ω*) is the power spectrum of the noise-corrupted signal *y(t) = s(t*) + *n(t*). Equivalently, since we can substitute {*F(ω*)}_*c*_ = *F(ω*) - {*F(ω*)}_*ac*_ for any *F(ω*), the optimality condition can be rewritten as:

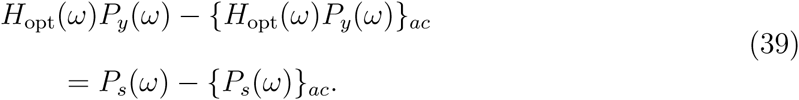

Divide both sides of Eq. (39) by 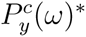, and then take the causal additive part {·}_*c*_ of both sides. The result is:

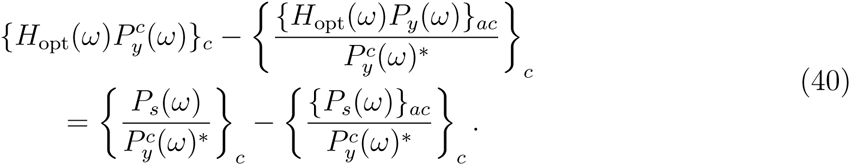

The second terms on both the left and right hand sides are the causal parts of a ratio between two anticausal functions. Since a ratio of anticausal functions is also anticausal, these terms are zero. On the left hand side the first term 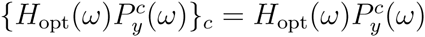, since *H*_opt_(*ω*) and 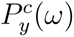 are causal, and hence their product is also causal. Making these simplifications, we can then solve for *H*_opt_(*ω*) as:

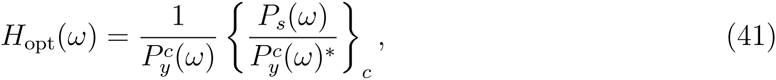

which is the optimal WK filter result shown as Eq. (9) in the main text.

## Appendix B: Linear response and noise filter analysis for a regulatory cascade

As an example of how our theory generalizes to control networks with multiple mediator species, we will consider the case where the feedback loop consists of a regulatory cascade. We will still explicitly single out a target species *R* and a mediator *P*, but now the signaling pathway which communic rates changes from *R* to *P* will be more complicated, consisting of a cascade of *N* species *Uj, j* = 1,…, *N*, with populations *u*_*j*_. The production of the *j*^*th*^ species will depend on the population of the *(j* - 1)^*th*^ species (with *j* = 0 corresponding to), and *P* will depend on the last member of the cascade, *U*_*N*_. In terms of Fourier-transformed fluctuations *δu*_*j*_, the dynamical equations for the pathway have the form:

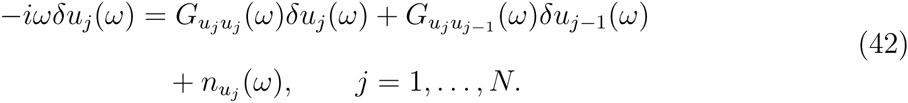

Thus the dynamics includes three parts: (i) the self-responses *G*_*u*_*j*___*u*_*j*__ which we can assume in the simplest case to be given by the inverse decay lifetimes of the species, 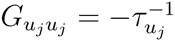; (ii) the cross-response terms *G*_*u*_*j*___*u*_*j-1*__ which describe how the *j*_*th*_ member of the cascade is related to the *(j* - 1)th member; (iii) the stochastic noise terms *n*_*uj*_. To complete the description of the feedback loop, we specify the equations for *R* and *P*:

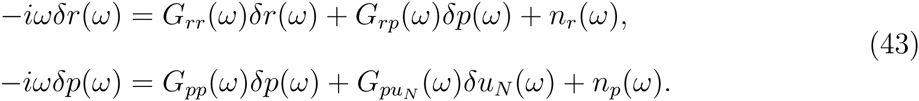

Instead of the simple cross-response *G*_*pr*_ from *R* to *P, P* is influenced by the final species of the *U*_*j*_ pathway through *G*_*pu*_*N*__.

The regulatory cascade system described by Eqs. (42)-(43) can in fact be simplified extensively, by solving for the dynamics of the mediator species *U*_*j*_ and substituting the results into Eq. (43). This yields equations for *R* and *P* which have the same form as in the two-species case in the main text, but with an effective cross-response function 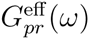 and noise term 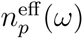,

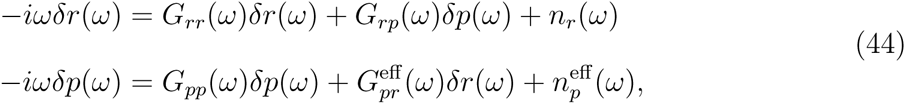

where:

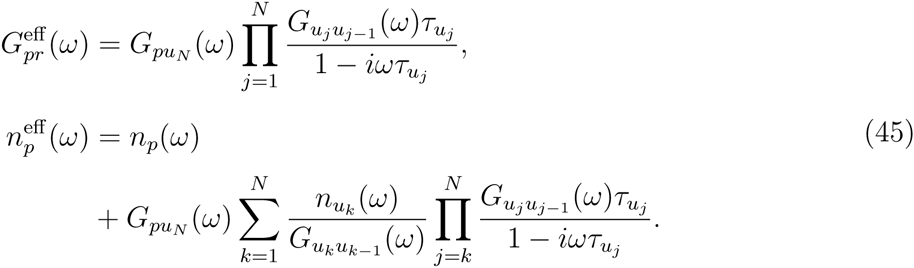

In this effective two-species reduction of the full system, all the stochastic effects of the mediators in the *U*_*i*_ pathway enter in as “extrinsic” noise contributions to 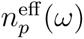. This is a particular example that shows how extrinsic noise encapsul rates the stochastic influence of all the species that are not explicitly specified in the dynamical equations.

The mapping of the two-species system onto the noise filter formalism, and the calculation of the optimal filter, can be carried out by the methods outlined in the main text. While this in general results in a more complicated problem than the simple example analyzed in the main text, in one scenario the noise filter optimization problem for the cascade is relatively straightforward: (i) we assume linear production functions 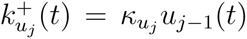 for all *U*_*j*_, so the cross-responses are constants in frequency space, *G*_*u*_*j*___*u*_*j-1*__(*ω*) = *κ*_*u*_*j*__. Similarly, the *P* production function is *κ*_*p*_*u*_*N*_(*t*), so *G*_*pu*_*N*__(*ω*) ≡ *κ*_*p*_. (ii) We assume the decay timescales of all the cascade species are negligible, *τ*_*u*_*j*__ ≪ *τ*_*r*_, so we can take the limits *τ*_*u*_*j*__ → 0 in Eq. (45). However, the products *κ*_*u*_*j*__*τ*_*u*_*j*__ remain finite for all *j*, since from the equilibrium conditions of the cascade (balance of production and destruction), they are related to ratios of the steady-state populations *ū*_*j*_:

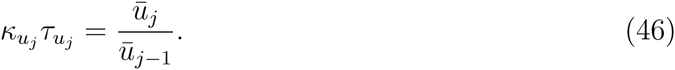

Hence rapid decay goes hand in hand with fast production. This is the same type of serial cascade analyzed in Ref. 19, where it was shown to maximize information transfer along the pathway. (iii) Finally, we assume that each species in the original, full description of the system is subject only to intrinsic noise, so the noise functions are given by:

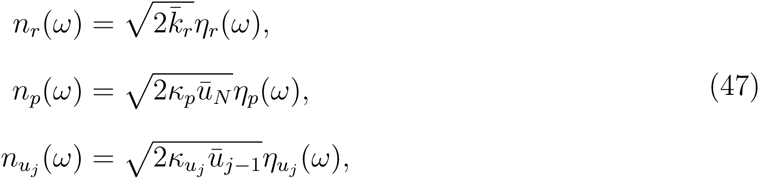

where the *η*_*α*_*(ω*) for different *α* are independent Fourier-transformed Gaussian white noise functions.

With these assumptions the effective cross-response and noise functions in Eq. (45) become:

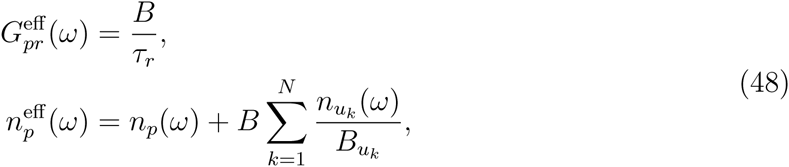

where the *P* burst ratio 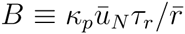 is analogous to *B* in the main text, i.e. the average number of *P* molecules produced per *R* during the time interval *τ*_*r*_. Similarly the burst ratio 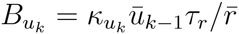 is the average number of *U*_*k*_ molecules produced per *R* during *τ*_*r*_.

The resulting signal and noise power spectra within the filter formalism are:

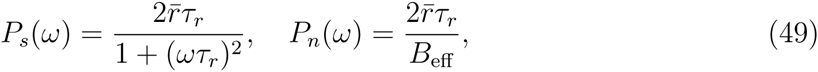

where

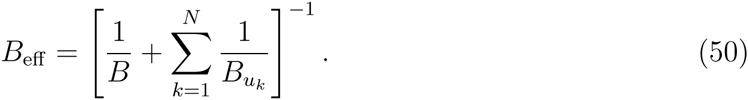

Since the power spectra in Eq. (49) have the same form as Eq. (12), with *B* replaced by *B*_*eff*_, all the subsequent optimality results are identical, but expressed in terms of the effective total burst ratio *B*_*eff*_ of the signaling pathway. This agrees with the effective burst ratio for the cascade derived by the information theory approach in Ref. 19, under the assumptions of rapid production/decay outlined above. Physically, this result implies that *B*_*eff*_ will be dominated by the smallest values among the *B* and *B*_*u*_*k*__. Hence, the efficiency of the noise filtration in the cascade is limited by the weakest links.

## Appendix C: Analytic limiting form of the generalized nonlinear feedback network

We will use the numerical optimization results described in the main text for the generalized nonlinear TetR feedback network (Eq. (22)) to derive a limiting form of the system that can be solved analytically. Since the optimization algorithm results in steep step-like functions *K*_*r*_*(p*) and Γ_*p*_*(p*) with thresholds at 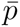, let us assume that optimal limit for these Hill functions looks like:

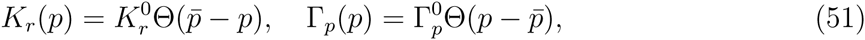

where the Heaviside step function Θ *(x*) = 0 for *x<*0 and *Θ (x*) = 1 for *x>*0. The plateau heights 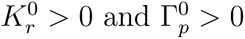 are assumed to be large, with a well defined ratio 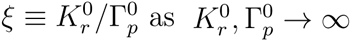. Since 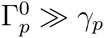 and thus Γ_*p*_*(p*) acts as the dominant protein degradation term, we will set *γ*_*p*_ = 0 for simplicity. (This has negligible effect on the resulting 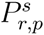, particularly since 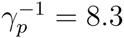 h was already the longest time scale in the system.)

Under these assumptions, we would like to find an analytical steady-state probability distribution 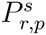 which satisfies *R*_*rp*_ = 0 from Eq. (27) for all *r, p≥*0. We cannot solve the system of equations directly, but we will introduce an ansatz for 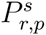 and verify that it is a solution to Eq. (27). The first part of the ansatz is trivial: we assume 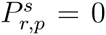 for 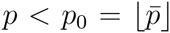. This satisfies *R*_*rp*_ = 0 for *p<p*_0_ exactly, regardless of the values of 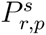 at *p ≥ p*_0_. To motivate the second part of the ansatz, which covers the *p ≥ p*_0_ region, we need some more information about the moments of the distribution. This can be gathered by defining the generating function,

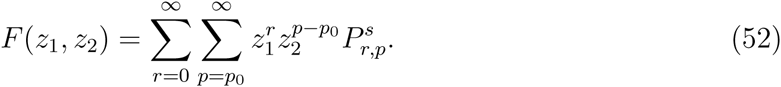

Summing the steady-state conditions *R*_*rp*_ = 0 in Eq. (27) over all *r*>0, *p≥p*0, we obtain an equation that can be expressed in terms of *F*:

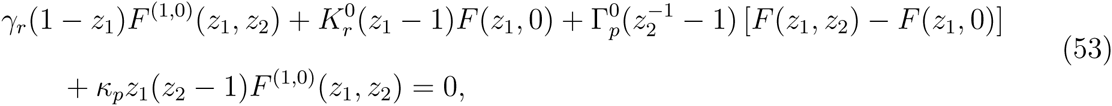

where 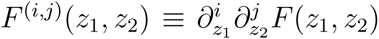. Taking the *z*_1_ derivative of Eq. (53), and evaluating the result at *z*_1_ = *z*1, *z*2 = 1, gives:

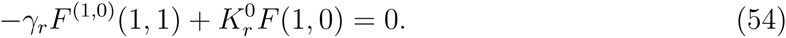

Similarly, differentiating Eq. (53) with respect to *z*_2_ yields:

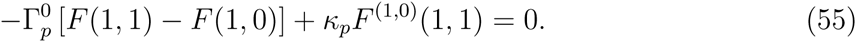

Using the fact that *F*(1, 1) = 1 from the normalization of 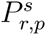, and *F*^(1,0)^(1,1) = 〈*r*〉, 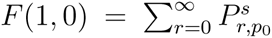 from the definition of the generating function in Eq. (52), we can use Eqs. (54) and (55) to find:

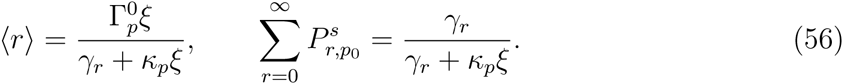

Thus we have an analytical expression for 〈*r*〉, one of the moments necessary for calculating the Fano factor. If we proceed to the next order of derivation, applying 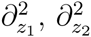, and *∂*_*z*_1__*∂*_*z*_2__ on Eq. (53) and evaluating at *z*_1_ = 1, *z*_2_ = 1, we can extract from these three equations the following moment relations:

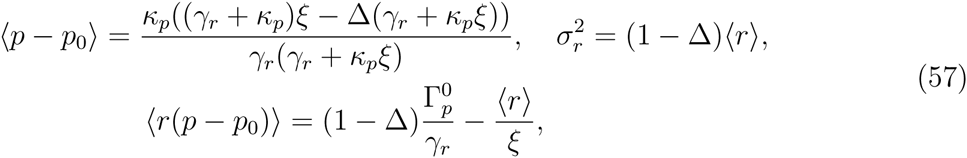

where ∆ is defined as

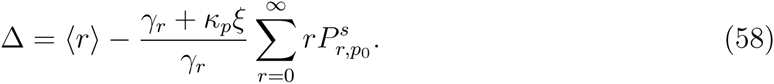

Thus the Fano factor 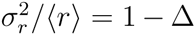, but unfortunately we do not have an explicit solution for ∆ from the generating function approach. (Higher order partial derivatives of Eq. (53) do not form a closed system of equations.) However, the moment relations in Eq. (57) will prove useful below.

From Eq. (56) we note that 〈*r*〉 → ∞ as 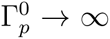, so the distribution is pushed toward larger *r* as the step functions become steeper, just as we saw in the numerical optimization (Fig. 6). In the large *r* limit, we can approximate 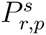, as a continuous function of *r* (though it remains discrete in *p*). Based on the numerical optimization results, we choose the following Gaussian ansatz for 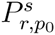, the first non-negligible *p* slice of the distribution:

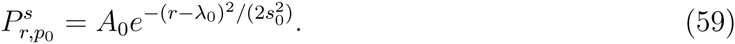

The parameters λ_0_ and *s*_0_ are to be determined, while *A*_0_ must be chosen to satisfy 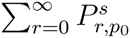 from Eq. (56). In the continuum, large limit we can approximate the sum as 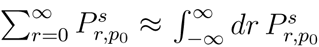 which implies that

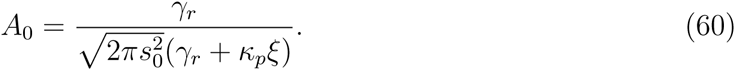

Similarly, Eq. (58) gives

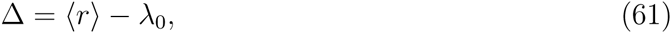

so finding λ_0_ is equivalent to finding ∆.

Let us now show that the ansatz of Eq. (59) yields a solution 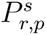 for *p≥p*_0_ that satisfies Eq. (27) in the large 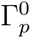 limit. Using Eq. (51) and the continuum approximation along the direction, we canewrite Eq. (27) for *p≥p*_0_ as

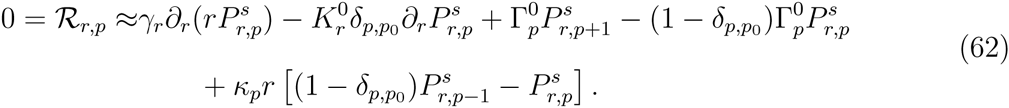

Plugging the ansatz for 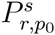 from Eq. (59) into Eq. (62) for *p = p*_0_, we can solve for 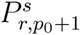

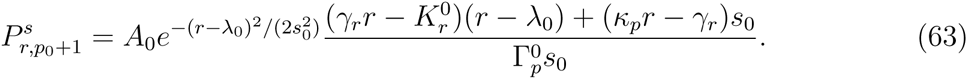

Similarly, once 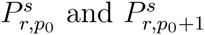 are known, Eq. (62) for *p* = *p*_0_ + 1 yields 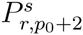

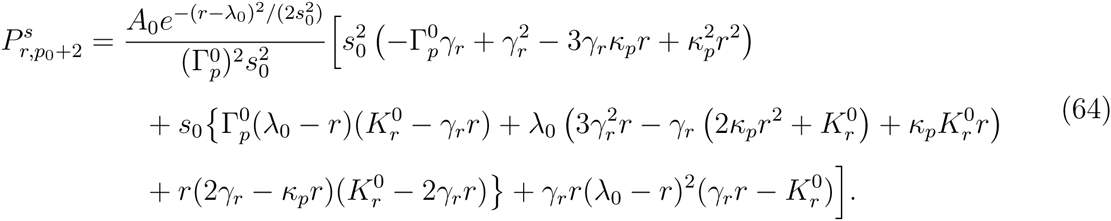

We can iterate this procedure, using Eq. (62) to generate analytical expressions for all 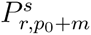, *m*> 0, which depend on the unknown parameters λ_0_ and *s*_0_. To solve for these parameters, let us first enforce the normalization condition,

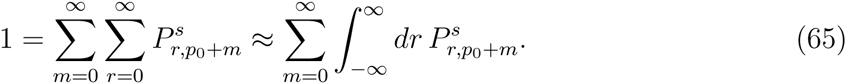

Though tedious, the integrals on the right-hand side of Eq. (65) can be explicitly carried out for each *m*, since 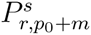 has the form of a Gaussian exp(-(*r* - λ_0_)^2^/(2*s*_0_)) times a polynomial in *r*. Since we are interested in the large 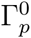 limit, we can Taylor expand the integrals up to first order in the small variable 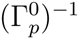, which gives the following result:

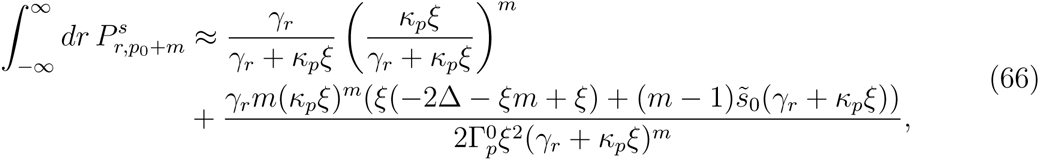

where 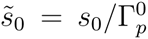, and we have used Eq. (61) to write λ_0_ = 〈*r*〉 - ∆, and Eq. (56) for 〈*r*〉. Plugging Eq. (66) into Eq. (65) and carrying out the sum over *m*, the normalization condition becomes

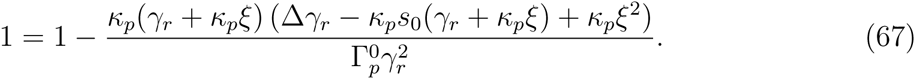

Thus the term of order 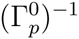 on the right must be zero, implying the following relation between 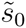 and ∆,

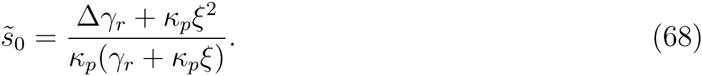

In order to complete the derivation and solve for ∆, we need to calculate the moment 〈*p-p*_0_〉,

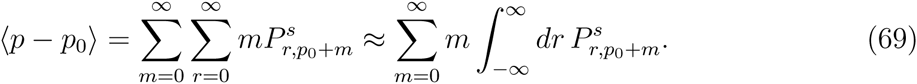

Plugging in Eq. (66) for the integral, we carry out the sum over *m* and simplify using Eq. (68), giving

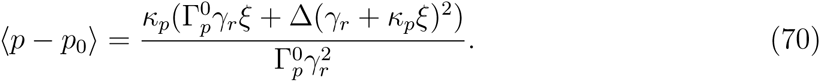

Setting this equal to the 〈*p-p*_0_〉 result from Eq. (57), we finally can solve for ∆, or equivalently the Fano factor 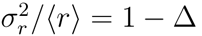

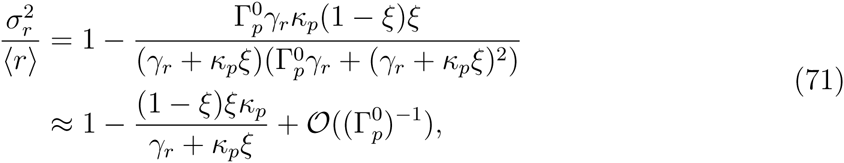

keeping the leading terms in the Taylor expansion for small 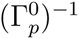. The Fano factor achieves a minimum value equal to the WK linear optimum,

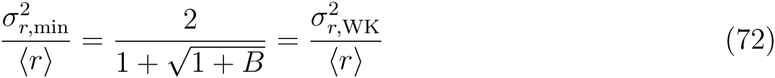

at 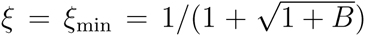, where *B = κ*_*p*_/*γ*_*r*_. Thus we see explicitly that nonlinear threshold regulation with *K*_*r*_(*p*) and Γ_*p*_*(p*) behaving like step functions can directly match (but not improve on) the efficiency of the optimal WK linear filter, so long as 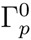 is large and the ratio of the step function heights assumes a particular value *ξ*_min_. Counter intuitively, this occurs despite the fact that the *p* copy numbers can be very small in our system, with a narrow range of fluctuations in which discreteness plays a major role.

## Appendix D: Optimality for the TetR gene network under extrinsic noise

In the frequency domain, we will model 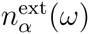, the extrinsic part of the noise associated with species *α* using,

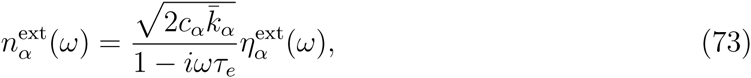

where *c*_*α*_ is a coefficient measuring the strength of the noise, and *η*^ext^(*ω*) is a Fourier-space Gaussian white noise function. Comparing to the definition of the intrinsic noise, 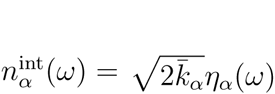, we see that *c*_*α*_ is the ratio of the extrinsic to intrinsic noise PSD for species *α* at *ω* = 0. The (1 - *iωτ*_*e*_)^−1^ factor acts as a cutoff that suppresses frequencies 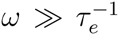. The total noise function for species *α* is the sum of intrinsic and extrinsic contributions, 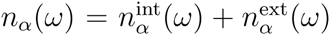. We will focus on how the addition of extrinsic noise affects the optimality conditions using the TetR yeast gene circuit example.

The calculation of *H*_opt_(*ω*) proceeds analogously to the no-extrinsic-noise procedure described in the main text. The power spectra of the signal and noise are,

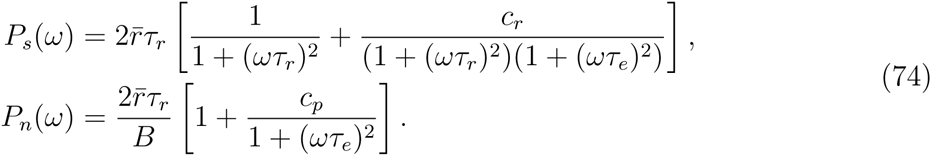

The first and second terms in the square brackets represent the intrinsic and extrinsic contributions respectively. The latter is parameterized by the coefficients *c*_*r*_ and *c*_*p*_, and the timescale *τ*_*e*_, which is assumed to be much larger than the dominant timescale, *τ*_*r*_, characterizing the *R* fluctuations. The signal plus noise power spectrum, *P*_*y*_(*ω*) = *P*_*s*_(*ω*) + *P*_*n*_(*ω*), can be rewritten as a causal decomposition in the following manner:

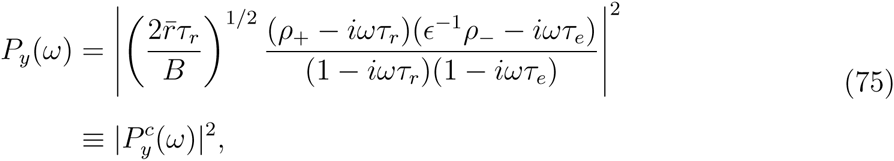

where ∊ ≡ *τ*_*r*_/*τ*_*e*_, and

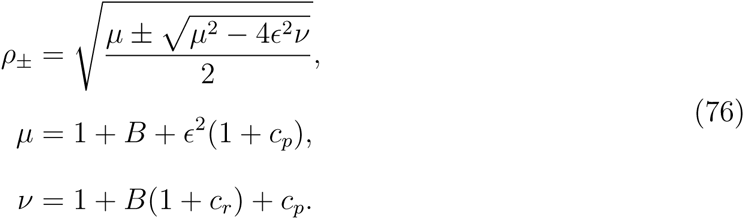

The expression 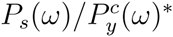 and its additive causal decomposition 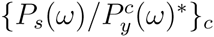 is given by:

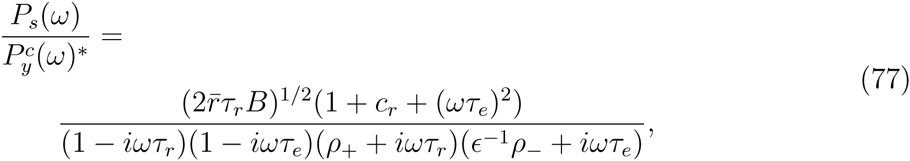

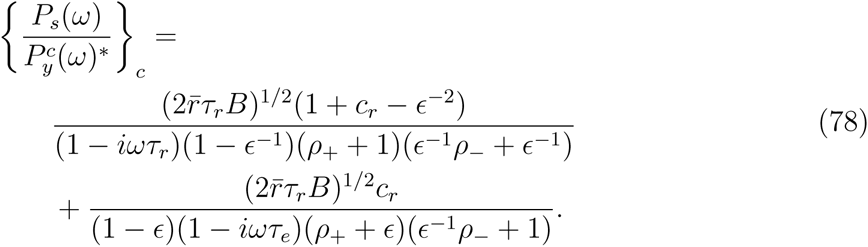

Using Eqs. (78) and (75) in Eq. (9), we obtain the form for the optimal filter function:

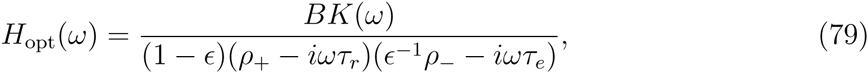

where

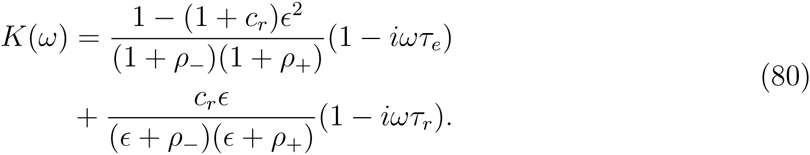

Since *∊* is presumed small, we will expand *H*_opt_ to lowest order in *∊*, giving the approximate expression:

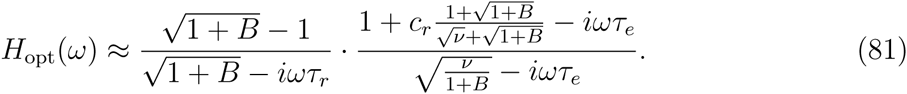

The first rational term is just the optimal filter result in the intrinsic-only case, Eq. (17), while the second term represents the modification needed to accommodate the extrinsic noise. As expected, the latter term approaches 1 when *c*_*r*_, *c*_*p*_ → 0, since *v* → 1 + *B* in this limit.

There is a different non-trivial scenario where the second term is equal to 1. If the noise magnitudes *c*_*r*_ and *c*_*p*_ are related such that,

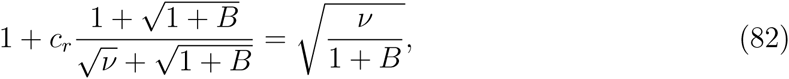

then the numerator and denominator exactly cancel each other out, removing the *τ*_*e*_ dependence from the optimal filter. Using the definition *v = 1 + B(1 + c*_*r*_) + *c*_*p*_, Eq. (82) can be simplified to yield the relation:

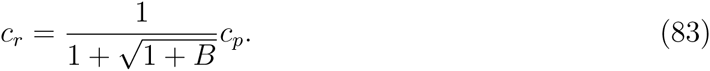

If this condition is satisfied, *H*_opt_(*ω*) is identical to the intrinsic-only optimal filter of Eq. (17) (to lowest order in ∊), and hence the approximate optimality is also achieved at the same feedback value, 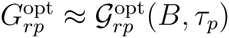.

Thus, the yeast gene circuit can still be fine-tuned to approach a WK optimal filter even in the presence of extrinsic noise. However, this tuning requires the relative strengths *c*_*r*_ and *c*_*p*_ of the *R* and *P* extrinsic noise to be related (at least approximately) by Eq. (83). The resulting minimal possible Fano factor 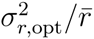 is:

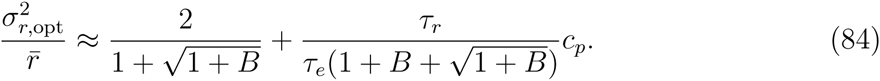

This is the intrinsic-only result of Eq. (21) in the main text plus an extrinsic noise contribution in the second term. Not surprisingly, with more total noise in the system, the standard deviation of the optimally filtered output increases. Since the second term is of the order *τ*_*r*_/*τ*_*e*_ it follows that the bigger the difference in time scales between the extrinsic noise (*τ*_*e*_) and the mRNA dynamics (*τ*_*r*_), the easier it is to filter out the extrinsic influence on the mRNA fluctuations. For *B* ≫ 1, the fundamental limit on the noise suppression still arises from the intrinsic term in 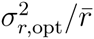, which scales like ~ *B*^−1/2^; the extrinsic contribution decays more rapidly, ~ *B*^−1^.

The blue curves in Fig. 7 show the linear theory predictions for 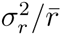 as a function of *A* in two cases: (i) *c*_*p*_ = 80, *c*_*r*_ = 23; (ii) *c*_*p*_ = 160, *c*_*r*_ = 46. The burst ratio *B* = 5, and *τ*_*e*_ is set equal to 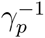, the longest time scale among the experimentally fitted parameters. For both these cases the noise strengths *c*_*p*_ and *c*_*r*_ satisfy the relation in Eq. (83), and hence it is possible to tune the system to approximately achieve WK optimality, just as in the intrinsic-only scenario. The noise magnitudes were chosen so that the system is noticeably perturbed by the extrinsic contribution. For example, if the signal *s(t*) is split into intrinsic and extrinsic parts *s*^int^*(t*) and *s*^ext^*(t*), the ratios of their respective standard deviations are 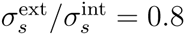 for case (i) and 1.6 for case (ii). The value of 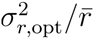 is marked by horizontal dashed lines, and the point *A = A*_opt_, where 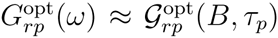 is satisfied, by a filled circle. In all cases the system approaches 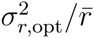 near *A = A*_opt_, verifying the optimality prediction.

As in the intrinsic-only scenario discussed in the main text, we can test the usefulness of the linear theory through Gillespie simulations (results shown as open squares and circles in Fig. 7), and reach a similar conclusion even in the presence of extrinsic noise. At large volumes, *V* = 10*V*_0_, the simulations converge to the linear theory, whereas for the more realistic volume *V* = *V*_0_ we see discrepancies due to nonlinearity and low copy numbers (*V*_0_ = 60 fL). Nevertheless, the Fano factor still reaches a minimum close to the predicted *A*_opt_ and 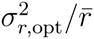 values.

## Graphical TOC Entry

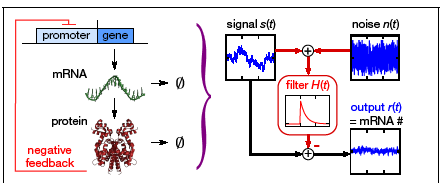

